# TAK1 is a key regulator of oncogenic signaling and differentiation blockade in rhabdomyosarcoma

**DOI:** 10.1101/2025.11.03.686399

**Authors:** Anh Tuan Vuong, Aniket S. Joshi, Anirban Roy, Kavya Mathukumalli, Phuong T. Ho, Raksha Bhat, Meiricris Tomaz da Silva, Tagari Samanta, Meghana V. Trivedi, Bin Guo, Benny A. Kaipparettu, Ashok Kumar

**Author notes:** **Corresponding author:** Ashok Kumar, Ph.D., Institute of Muscle Biology and Cachexia, Department of Pharmacological and Pharmaceutical Sciences, Health Building 2, Room 5012, College of Pharmacy, University of Houston, Houston, TX 77204-1217.

## Abstract

Rhabdomyosarcoma (RMS) is a malignant soft tissue sarcoma with a skeletal muscle phenotype, accounting for approximately 50% of all pediatric soft tissue sarcomas and 8% of all childhood cancers. Although RMS cells express myogenic regulatory factors, they fail to undergo terminal differentiation into mature muscle cells. Transforming growth factor β-activated kinase 1 (TAK1) is a major signaling protein that activates multiple intracellular pathways in response to growth factors, cytokines, and microbial products. Emerging evidence suggests that TAK1 is also an important regulator of self-renewal, proliferation, and differentiation of muscle progenitor cells. However, the role and mechanisms of action of TAK1 in RMS remain completely unknown. In this study, we demonstrate that TAK1 expression and activity are markedly elevated in a panel of RMS cell lines and in patient tumor specimens. Reverse phase protein array (RPPA) analyses revealed that TAK1 regulates the expression and activity of many molecules involved in cell cycle control, cell proliferation, and oncogenic signaling. Genetic knockdown or pharmacological inhibition of TAK1 suppresses RMS cell proliferation, migration, and invasiveness, while also promoting terminal myogenic differentiation. TAK1 inhibits differentiation in RMS, at least in part, through up-regulating YAP1 signaling. Our results also demonstrate that inducible knockdown of TAK1 in human RMS xenografts retards tumor growth and enhances myogenic differentiation *in vivo*. Collectively, these findings uncover a previously unrecognized role for TAK1 in RMS growth and differentiation, and suggest that TAK1 can be a potential therapeutic target for the treatment of RMS.

## Introduction

Rhabdomyosarcoma (RMS) is the most common soft tissue sarcoma in children that arises in or near skeletal muscle beds. It poses significant challenges due to its aggressive nature and high metastatic potential (1, 2). RMS includes two main histological subtypes, namely embryonal RMS (ERMS) and alveolar RMS (ARMS) which differ in their molecular drivers. ARMS are driven mainly by an oncogenic chromosomal translocation between paired box 3 (*PAX3)* or PAX7 and the Forkhead transcription factor (FOXO1) and are designated as fusion-positive RMS. In contrast, ERMS is devoid of any fusion gene but often harbor mutations in p53 (TP53) and RAS, amplification of *CDK4*, and upregulation of *MYCN* (3–5). Regardless of their cell of origin (muscle or non-myogenic precursor), RMS tumors exhibit myogenic characteristics, such as the expression of myogenic regulatory factors (MRFs) and skeletal muscle structural proteins (6–8). However, despite expressing various MRFs, RMS cells remain proliferative and fail to undergo terminal differentiation into mature skeletal muscle. This blockade in differentiation suggests that therapeutic agents capable of inhibiting proliferation while promoting myogenic differentiation may offer a promising strategy for RMS treatment (5, 9).

Similar to many other cancer types, RMS exhibits defects in cell cycle checkpoints, growth factor signaling, and tumor suppressor pathways, which contribute to unrestricted proliferation and impaired differentiation (8, 10). Several studies have shown that key oncogenic pathways activated by fibroblast growth factor (FGF) and insulin-like growth factor (IGF), including RAS-RAF-MAPK and PI3K-AKT-mTOR, are dysregulated in RMS tumors (2, 9, 10). The nuclear factor-kappa B (NF-κB) pathway has also been implicated in promoting proliferation and inhibiting terminal differentiation of RMS cells (8, 11, 12). Accumulating evidence further indicates that multiple developmental signaling pathways, such as Notch, Wnt, Hippo-YAP1, and Hedgehog (Hh), are aberrantly activated in RMS. These pathways normally regulate the balance between self-renewal and differentiation in muscle progenitor cells, and their dysregulation in RMS is thought to shift the balance towards uncontrolled proliferation and a block in terminal differentiation (13–24). Despite these observations, the upstream signaling events that regulate the activation of various intracellular signaling pathways and the specific role each pathway plays in the progression of RMS remain incompletely understood.

Transforming growth factor β-activated kinase 1 (TAK1, gene name: MAP3K7) is a key signaling protein that mediates the activation of multiple downstream pathways, including MAPK cascades and the IκB kinase β (IKKβ)-NF-κB pathway, in response to various growth factors and pro-inflammatory cytokines (25–28). TAK1 functions as a part of a heterotrimeric complex with TAB1 and either TAB2 or TAB3, which interact with the N- or C-terminus of TAK1, respectively (29). Activation of the TAK1 signalosome is triggered by K63-linked polyubiquitination, catalyzed by the E2 conjugating enzyme complex UBC13/UEV1A together with the RING-type E3 ligases TRAF2 or TRAF6, in response to inflammatory or stress stimuli. Specifically, K63-linked polyubiquitination at lysine 158 (K158) of TAK1 serves as a crucial modification that facilitates its activation (30, 31). Following ubiquitination, TAK1 undergoes autophosphorylation at threonine 187 (Thr187) within its activation loop and may also phosphorylate additional regulatory residues, including Thr184 and Ser192 (30, 31). TAK1 plays an essential role in the regulation of cell proliferation, survival, and differentiation, and its aberrant activation has been implicated in the progression and metastasis of several cancers, including osteosarcoma, as well as esophageal, thyroid, gastric, breast, and ovarian cancers (32–35).

Accumulating evidence also suggests that TAK1 is an important regulator of skeletal muscle development and homeostasis (36, 37). We have previously reported that TAK1 and its upstream adaptor TRAF6 are essential for the self-renewal of muscle progenitor cells in skeletal muscle of adult mice. Targeted deletion of TAK1 impairs muscle regeneration by reducing survival and inducing premature differentiation of muscle progenitor cells (38, 39). Additionally, TAK1 promotes the differentiation of cultured myoblasts into multinucleated myotubes (40, 41). However, the regulation of TAK1 activity in RMS and its precise role in RMS tumor biology remains unexplored.

In this study, we investigated the role of TAK1 in RMS growth and progression using complementary in vitro and in vivo approaches. Our findings reveal that both the expression and phosphorylation of TAK1 are markedly elevated in ARMS and ERMS cell lines, as well as in patient tumor samples. Genetic knockdown or pharmacological inhibition of TAK1 significantly reduced RMS cell proliferation, migration, and invasiveness. Notably, TAK1 inhibition also promoted terminal myogenic differentiation in RMS cells. Mechanistically, TAK1 appears to suppress myogenic differentiation, at least in part, by upregulating the pro-oncogenic YAP1 protein. Finally, inducible TAK1 knockdown in RMS xenografts impaired tumor growth, decreased YAP1 levels, and induced cellular differentiation.

## Results

### TAK1 expression and activity is increased in RMS cell lines and in patient tumor specimens

We first analyzed how the gene expression of various components of TAK1 signalosome is affected in human RMS samples. Analysis of a publicly available microarray dataset (GSE141690) that includes 16 normal muscle and 66 RMS samples revealed that the mRNA levels of TAK1, TAB1, TAB3, and TRAF6, but not TAB2, were significantly upregulated in RMS samples compared to normal muscle (**Fig. 1A**). We also analyzed another publicly available RNA-seq dataset (GSE108022) that includes normal muscle (n=5), fusion-positive RMS (FP-RMS, n=28) and fusion-negative RMS (FN-RMS, n=51) samples. This analysis also showed that the mRNA levels of TAK1, TAB1, TAB3 and TRAF6 are significantly up-regulated in both FP-RMS and FN-RMS compared to normal muscle (**Fig. 1B**). We next investigated how the activity and levels of TAK1 are affected in cultured ERMS (i.e. RD, HTB8, and RH36) and ARMS (i.e. RH30 and RH41) cell lines compared with human myoblasts (HM). Phosphorylation of Thr184 and Thr187 residues within the TAK1 activation loop is essential for TAK1 kinase activity (42, 43). Overall, both phosphorylated and total TAK1 protein levels were elevated in ERMS and ARMS cell lines compared with HM cells. The levels of TAK1-associated adaptor proteins (TAB1, TAB2, TAB3) and the E3 ligase TRAF6 were variably increased across RMS cell lines, with most showing higher levels than HM (**Fig. 1C-F**). These findings suggest that TAK1 signaling components are broadly upregulated and activated in RMS subtypes.

**FIGURE 1.**
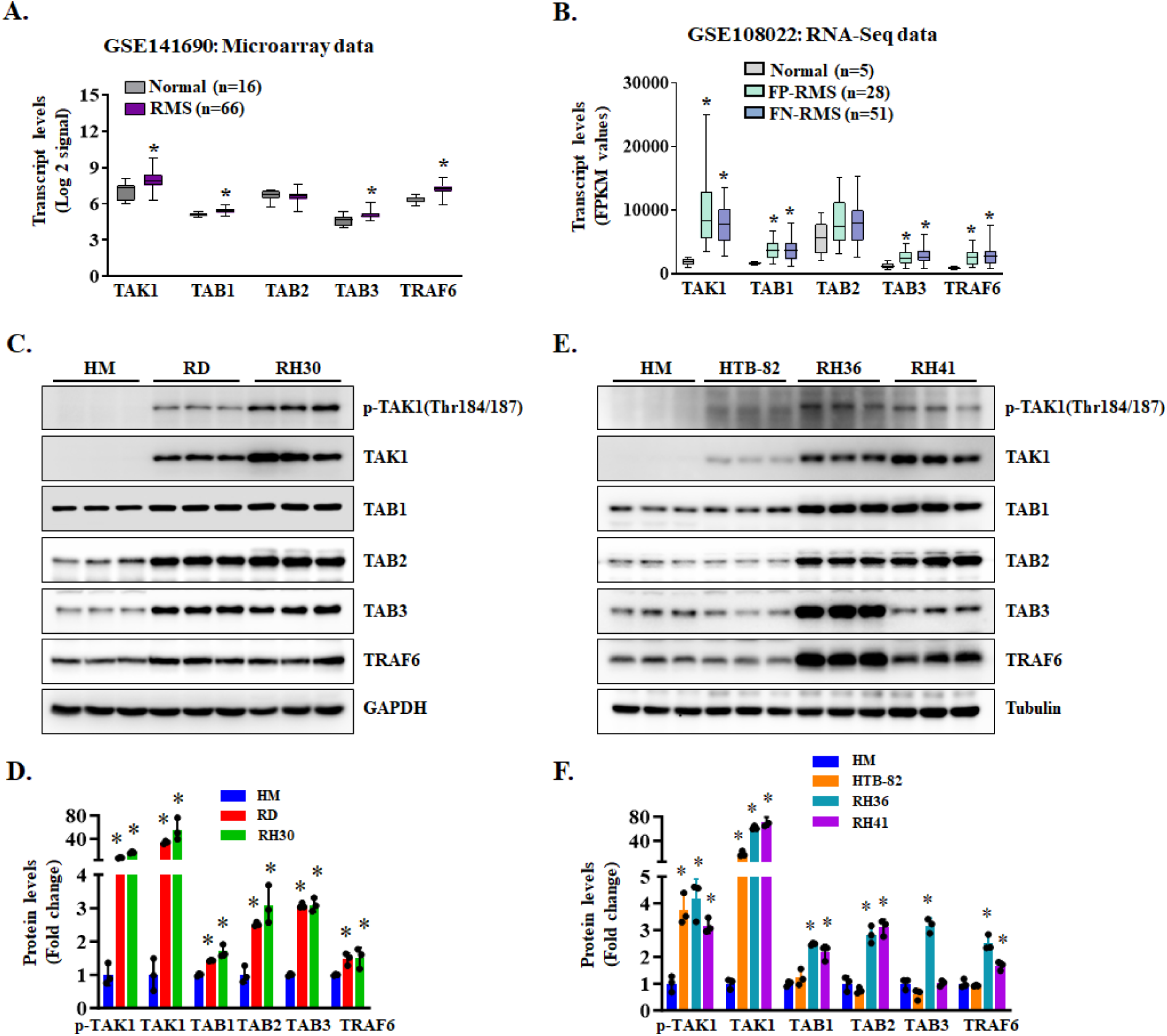
Expression and activation of TAK1 in RMS cell lines and tumor samples. **(A)** Relative mRNA levels of TAK1 obtained from analysis of GSE141690 microarray dataset containing 66 RMS and 16 normal muscle samples. *p < 0.05, value significantly different from normal muscle samples by unpaired two-tailed t-test. **(B)** Relative mRNA levels of TAK1 obtained from analysis of RNA-Seq dataset (GSE108022) containing 5 normal muscles, 28 FP-RMS and 51 FN-RMS samples. *p < 0.05, values significantly different from normal muscle samples by unpaired two-tailed t-test. **(C)** Representative immunoblots, and **(D)** densitometry analysis showing levels of p-TAK1 (Thr184/187), TAK1, TAB1, TAB2, TAB3, TRAF6, and unrelated protein GAPDH in human myoblasts (HM) and RD and RH30 cell lines. **(E)** Immunoblots and **(F)** densitometry analysis demonstrating the levels of p-TAK1, TAK1, TAB1, TAB2, TAB3, TRAF6, and unrelated protein tubulin in HM, HTB-82, RH36, and RH41 cells. N = 3 biological replicates in each group. Data presented as mean ± SD. *p<0.05, values significantly different from HM using unpaired two-tailed t-test.

### Silencing of TAK1 inhibits the proliferation of RMS cells

To understand the role of TAK1 in RMS, we first generated lentiviral constructs expressing a scrambled shRNA or TAK1 shRNA targeting two different regions of human TAK1 mRNA. RD cells were transduced with lentivirus expressing scrambled (control) or TAK1 shRNA # 1 or 2 for 48h followed by performing western blot for TAK1 protein. As shown in **Fig. 2A**, expression of either of the two TAK1 shRNA reduced the levels of TAK1. We next performed RNA-Seq analysis of scrambled- or TAK1-shRNA expressing RD cells in triplicates. Differentially expressed genes (DEGs) were identified using the threshold of |Log2FC| > 0.25 and adjusted *p*-value < 0.05. This analysis revealed that compared to control cells, 1508 genes were significantly (FDR < 0.05) upregulated whereas 1757 genes were significantly downregulated in TAK1 knockdown RD cells (**Fig. S1A**). Pathway enrichment analysis of DEGs showed that downregulated genes were associated with the processes of cell cycle, mitotic cell cycle, cellular response to stress, signaling by TGFβ family members, and mesenchyme development whereas upregulated genes were associated with cilium organization, muscle structure development, actin filament-based process, muscle cell differentiation, and regulation of cytoskeletal organization (**Fig. 2B**). Heatmap analysis of DEGs further showed that knockdown of TAK1 reduced the expression of various molecules involved in cell cycle regulation and oncogenic signaling (**Fig. 2C**).

**FIGURE 2.**
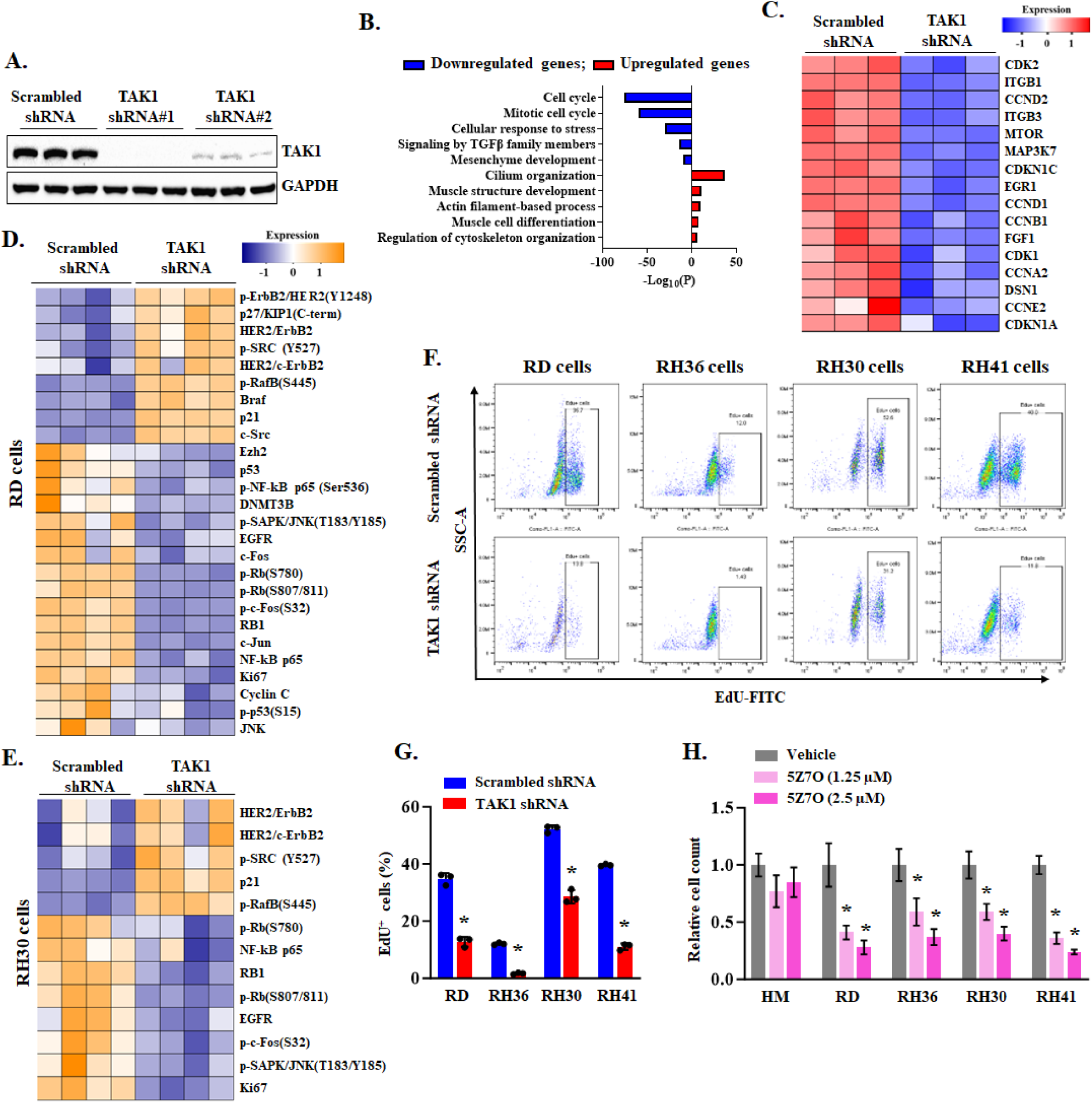
Role of TAK1 in the proliferation of RMS cells. **(A)** Representative immunoblots presented here show levels of TAK1 protein in RD cells expressing scrambled shRNA or two TAK1 shRNA. **(B)** Gene ontology (GO) biological processes associated with down-regulated and up-regulated genes in TAK1 knockdown RD cells. **(C)** Heatmap generated from RNA-Seq dataset showing regulation of selected genes involved in cell cycle regulation and proliferation in control and TAK1 knockdown RD cultures. Heatmaps generated from RPPA analysis showing differences in phosphorylated or total levels of various cell cycle-regulators and proto-oncogenes, oncogenes and tumor suppressors in TAK1 shRNA expressing **(D)** RD and **(E)** RH30 cells compared to their corresponding control cells expressing scrambled shRNA. **(F)** Representative scatter plots of FACS-based analysis demonstrate the EdU^+^ cells amongst all cells in control and TAK1 knockdown RD, RH36, RH30 and RH41 cells. **(G)** Quantification of proportion of EdU^+^ cells in control and TAK1 knockdown RD, RH36, RH30 and RH41 cells measured by FACS analysis. n = 3 (biological replicates) in each group. **(H)** Effect of indicated concentrations of 5Z7O on the proliferation of HM, RD, RH36, RH30, and RH41cells assessed by plating 1,000 cells per well in a 96-well plate. The graph illustrates the relative cell count on day 5 after treatment with indicated concentrations of 5Z7O. Data presented as mean ± SD. *p<0.05, values significantly different from corresponding cells treated with vehicle alone by unpaired two-tailed t-test.

We next performed antibody-based reverse-phase protein array (RPPA) that quantitatively analyzes around 260 cancer-related proteins and their activation states (44–46). We found that there were 30 common upregulated and 36 common downregulated proteins in RD and RH30 cells (**Fig. S1B, C**). Hallmark gene set enrichment analysis of downregulated proteins in RPPA dataset showed that E2F targets, PI3K-Akt-mTOR signaling, G2M checkpoint, glycolysis, Wnt β-catenin, and epithelial-mesenchymal transition were common downregulated pathways in TAK1 knockdown RD and RH30 cells (**Fig. S1D**). Heatmap analysis of RPPA dataset also showed that silencing of TAK1 in RD or RH30 cells significantly affected the total levels or phosphorylation of various cell cycle-regulators, proto-oncogenes, oncogenes and tumor suppressors (**Fig. 2D, 2E**). For instance, TAK1 silencing led to decreased phosphorylation of the retinoblastoma (Rb) protein and reduced levels of the proliferation marker Ki67, while upregulating the cell cycle inhibitor p21. TAK1 is an upstream kinase in signaling cascades that lead to the activation of MAPKs, nuclear factor-kappa B (NF-κB), and a few other pathways (26, 27). RPPA analysis showed that phosphorylation of JNKs and their downstream targets, such as c-Jun and c-Fos, and phosphorylation of NF-κB subunit p65 were significantly reduced in TAK1 knockdown RD or RH30 cells compared to their corresponding controls (**Fig. 2D, 2E**). We also performed independent western blot analysis to measure the phosphorylation of MAPKs. Consistent with RPPA, the phosphorylation of JNK1/2 and p38 MAPK was significantly reduced in TAK1 knockdown RD cells whereas the phosphorylation of ERK1/2 and JNK1/2 was significantly reduced in TAK1 knockdown RH30 cells compared to their corresponding controls (**Fig. S2**).

We next examined the impact of TAK1 knockdown on the proliferation of RMS cell lines. ERMS (RD, RH36) and ARMS (RH30, RH41) cells were transduced with lentiviral particles expressing either scrambled shRNA or TAK1-targeting shRNA, and cell proliferation was assessed on day 5 using an EdU (5-ethynyl-2′-deoxyuridine) incorporation assay followed by FACS analysis. The proportion of EdU□ nuclei was significantly reduced in TAK1 shRNA-expressing RD, RH36, RH30, and RH41 cells compared with their respective scrambled shRNA controls (**Fig. 2F, G**). To further evaluate the role of TAK1, we analyzed the effect of its pharmacological inhibition using 5Z-7-oxozeaenol (5Z7O), a potent and specific TAK1 inhibitor (39, 47). Primary human myoblasts (HM) and RMS cells (RD, RH36, RH30, RH41) were treated with either vehicle or different concentrations of 5Z7O, and cellular proliferation was measured by cell counting on day 5. Treatment with 5Z7O significantly reduced cell count in a dose-dependent manner across multiple RMS cell lines. In contrast, treatment with 5Z7O did not have any significant effect on the number of HM (**Fig. 2H**). Together, these results indicate that both genetic and pharmacological inhibition of TAK1 suppress RMS cell proliferation in ERMS and ARMS subtypes.

### TAK1 knockdown diminishes the survival of RMS cells

To determine whether TAK1 regulates the viability of RMS cell lines, RD, RH36, RH30, and RH41 cells were transduced with lentiviral particles expressing either scrambled control or TAK1-targeting shRNA. After 24 hours, cells were plated at equal densities, and apoptosis was assessed 96 hours later using Annexin V and propidium iodide (PI) staining followed by flow cytometry. TAK1 knockdown led to a modest but statistically significant increase in Annexin V cells in ERMS lines RD (∼10%) and RH36 (∼8%) compared with controls. The effect was more pronounced in ARMS lines, with approximately 18% of RH30 and 12% of RH41 cells staining positive for Annexin V, indicating greater sensitivity to TAK1 depletion (**Fig. 3A, B**). In parallel, an MTT assay was performed to assess cell viability based on mitochondrial metabolic activity. TAK1 knockdown significantly reduced MTT conversion in RD, RH36, RH30, and RH41 cells relative to controls (**Fig. 3C**), consistent with decreased proliferation and survival. To examine long-term effects, a clonogenic assay was conducted in ERMS (RD, RH36) and ARMS (RH30, RH41) cells. TAK1 knockdown markedly reduced clonogenic potential across all RMS cell lines tested (**Fig. 3D, E**). Similarly, treatment with the TAK1 inhibitor 5Z7O suppressed clonogenic potential in a dose-dependent manner across all RMS cell lines tested (**Fig. 3F, G**). Collectively, these results suggest that TAK1 is essential for maintaining the proliferative capacity, metabolic activity, and survival of both ERMS and ARMS cell lines.

**FIGURE 3.**
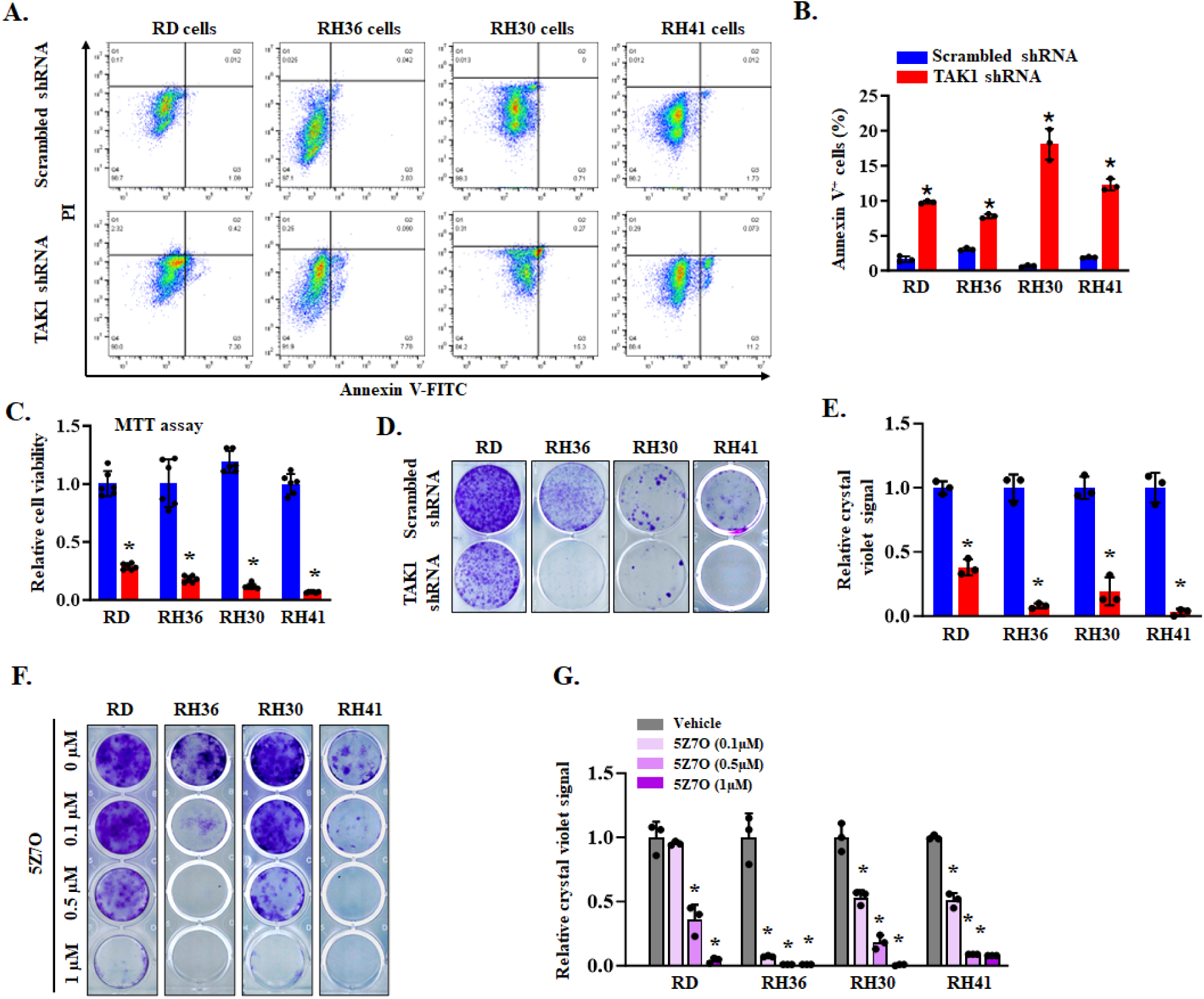
Silencing of TAK1 reduces the viability of RMS cells. **(A)** Representative scatter plots of FACS-based analysis demonstrate the Annexin V-positive cells amongst control and TAK1 knockdown RD, RH36, RH30 and RH41 cells. **(B)** Quantification of Annexin V^+^ cells in control and TAK1 knockdown RD, RH36, RH30 and RH41 cells measured by FACS analysis. n = 3 biological replicates in each group. **(C)** MTT assay demonstrating relative cell viability in control and TAK1 knockdown RD, RH36, RH30, and RH41 cells. n= 6 biological replicates in each group. **(D)** Representative images of clonogenic assay and **(E)** quantification of crystal violet dye in each well of control and TAK1 knockdown RD, RH36, RH30, and RH41 cells. n= 3 biological replicates in each group. **(F)** Representative images of clonogenic assay of RD, RH36, RH30, and RH41 cells after treatment with indicated concentrations of 5Z7O. **(G)** Quantification of crystal violet dye signal in vehicle and 5Z7O-treated cultures. n= 3 biological replicates in each group. Data are presented as mean ± SD. *p<0.05, values significantly different from corresponding RD, RH36, RH30, and RH41 cells expressing scrambled shRNA or treated with vehicle alone by unpaired two-tailed t-test.

### TAK1 promotes epithelial-mesenchymal transition (EMT) in RMS cells

EMT is a dynamic process characterized by a phenotypic and functional shift from an epithelial to a mesenchymal state, playing a critical role in tumor growth, invasion, and metastasis (48). Although RMS arises from mesenchymal tissue rather than epithelial cells, EMT-related transcriptional programs remain active and contribute to RMS progression by enhancing cellular motility and metastatic potential (49). EMT is regulated by multiple signaling pathways, including TGF-β, Wnt, and PI3K–AKT, and transcription factors such as Snail, Twist, and ZEB1 (50). Transcriptomic analysis of RNA-Seq data revealed that TAK1 knockdown in RD cells led to a significant downregulation of genes associated with EMT, including components of the TGF-β signaling pathway (**Fig. 4A**). Furthermore, RPPA dataset analysis showed that EMT was a common pathway downregulated in both RD and RH30 cells following TAK1 knockdown (**Fig. S1D**). Heatmap analysis of RPPA dataset also showed reduced phosphorylation and/or total levels of several EMT-related proteins, such as β-catenin, Wnt5a/b, ZEB1, and Slug, in both RD and RH30 cells following TAK1 silencing (**Fig. 4B, C**).

**FIGURE 4.**
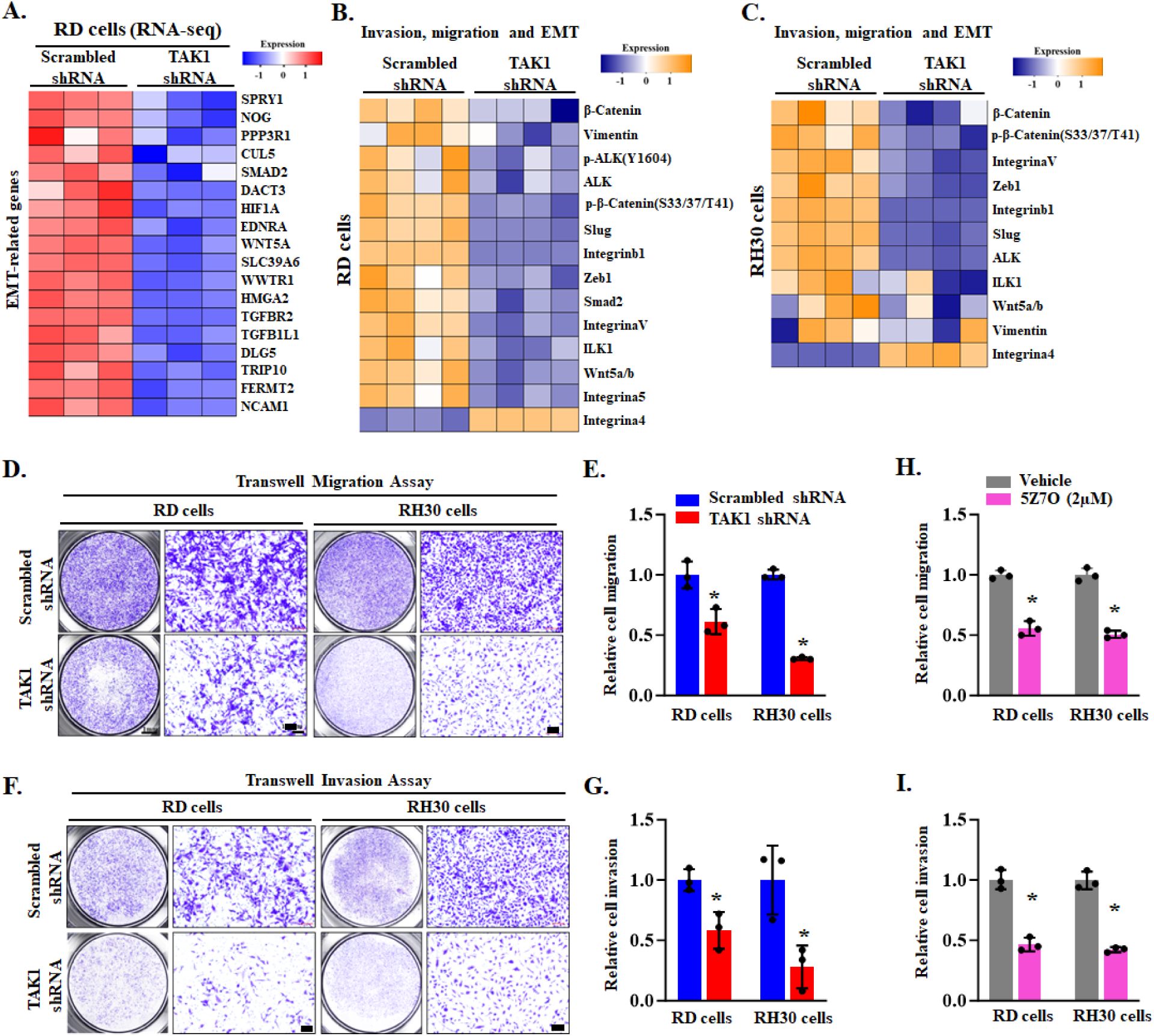
Silencing of TAK1 inhibits EMT signature and migration and invasion of RMS cells. **(A)** Heatmap generated from RNA-seq dataset showing expression of various EMT-related molecules in control and TAK1 knockdown RD cultures. Heatmaps generated from RPPA analysis showing differences in phosphorylated or total levels of various proteins involved in the regulation of EMT, migration, and invasion in control and TAK1 knockdown **(B)** RD and **(C)** RH30 cells. **(D)** Representative images of control and TAK1 knockdown RD and RH30 cells in transwell migration assay. Scale bars, 50 μm. **(E)** Quantification of the relative migration of control and TAK1 knockdown RD and RH30 cells. **(F)** Representative images of control and TAK1 knockdown RD and RH30 cells in transwell invasion assay. Scale bars, 50 μm. **(G)** Quantification of the relative invasive capacity of control and TAK1 knockdown RD and RH30 cells. n = 3 biological replicates in each group. Data are presented as mean ± SD. *p<0.05, values significantly different from corresponding RD or RH30 cells expressing scrambled shRNA by unpaired two-tailed t-test. Quantification of the effect of 5Z7O on relative **(H)** migration and **(I)** invasive capacity of RD and RH30 cells. n = 3 biological replicates in each group. Data are presented as mean ± SD. *p<0.05, values significantly different from corresponding RD or RH30 cells treated with vehicle alone by unpaired two-tailed t-test.

Since EMT enhances tumor cell migration and invasion, we next assessed the impact of TAK1 depletion on these cellular behaviors in representative ERMS (RD) and ARMS (RH30) cell lines. Transwell assays were performed within 24 hours of plating, a time point when TAK1 knockdown does not significantly affect RMS cell proliferation or viability. TAK1 silencing significantly reduced the migratory capacity of both RD and RH30 cells in transwell assays (**Fig. 4D, E**). Similarly, in matrigel-coated transwell assays, TAK1 knockdown markedly decreased the invasive potential of both cell lines (**Fig. 4F, G**). Parallel experiments using the TAK1 inhibitor 5Z-7-oxozeaenol (5Z7O) produced similar results, with significant reductions in both migration and invasion of RD and RH30 cells (**Fig. 4H, I**). Collectively, these results indicate that TAK1 promotes EMT-associated gene expression and enhances the migratory and invasive capacities of RMS cells.

### TAK1 knockdown augments the expression of muscle differentiation markers in RMS cells

While RMS cells express various MRFs, they fail to undergo terminal differentiation into myotubes. Pathway analysis of DEGs in control and TAK1 knockdown RD cells in RNA-Seq experiment revealed that silencing of TAK1 upregulates processes related to muscle cell differentiation and muscle structure development (**Fig. 2B**). Further analysis of DEGs showed that knockdown of TAK1 significantly increased mRNA levels of muscle differentiation markers (e.g., Myogenin, Myomaker, MyoD, HDAC4, HDAC9 and MYH3) in RD cells (**Fig. 5A**). To validate these findings, we examined the effect of TAK1 knockdown on the expression of muscle differentiation markers in RMS cell lines. RD, RH36, RH30, and RH41 cells were transduced with lentiviral particles expressing scrambled shRNA (control) or TAK1-targeting shRNA. After 48 h, the cells were plated at equal density in growth medium, and 72 h later, the expression of myosin heavy chain (MyHC) was assessed by performing immunohistochemistry and western blot. Remarkably, TAK1 knockdown led to a marked increase in MyHC^+^ cells across all RMS cell lines tested, with the most pronounced effect observed in RD cells (**Fig. 5B, C**). Furthermore, western blot analysis showed that the levels of MyHC and another muscle differentiation marker myogenin were considerably increased in TAK1 knockdown RD, RH36, RH30, and RH41 cells compared to their corresponding control cells (**Fig. 5D, E**). In a parallel experiment, we studied myogenic differentiation by measuring the activity of skeletal α-actin (SK) promoter reporter construct. Control and TAK1 knockdown RD or RH30 cells were transfected with empty vector (i.e., pGL4) or SK-Luc plasmid along with Renilla luciferase expressing plasmid. After 3 days, the cells were lysed and the luciferase activity in cell lysates was measured. Results showed that the activity of skeletal α-actin promoter was significantly higher in TAK1 knockdown RD or RH30 cells compared to their corresponding controls (**Fig. 5F**).

**FIGURE 5.**
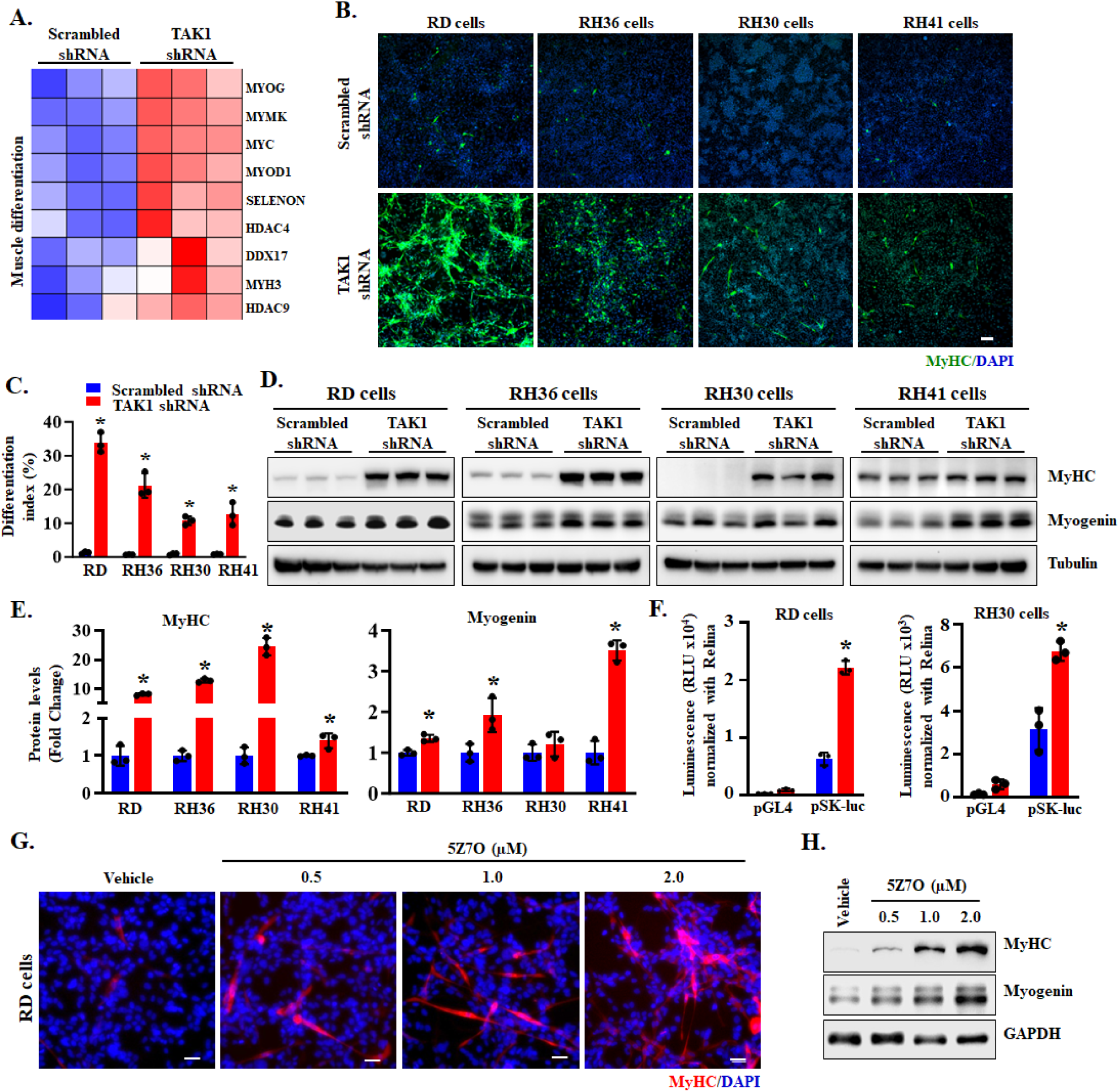
Silencing of TAK1 induces myogenic differentiation in RMS cells. **(A)** Heatmap generated from RNA-Seq dataset showing regulation of selected genes involved in muscle differentiation in scrambled shRNA and TAK1 knockdown RD cultures. **(B)** Representative images of control and TAK1 knockdown RD, RH36, RH30, and RH41 cells after immunostaining for myosin heavy chain (MyHC) protein. Nuclei were visualized by staining with DAPI. Scale bar: 100 µm. **(C)** Quantification of differentiation index in control and TAK1 knockdown RD, RH36, RH30, and RH41 cultures. **(D)** Immunoblots presented here demonstrate the levels of MyHC, myogenin, and unrelated protein tubulin in control and TAK1 knockdown RD, RH36, RH30, and RH41 cells. **(E)** Quantification of relative MyHC and myogenin protein levels between control and TAK1 knockdown RD, RH36, RH30 and RH41 cells. **(F)** Quantification of skeletal α actin (SK) reporter (luciferase, Luc) activity in control and TAK1 knockdown RD and RH30 cells. n = 3 biological replicates in each group. Data are presented as mean ± SD. *p<0.05, values significantly different from corresponding cells expressing scrambled shRNA by unpaired two-tailed t-test. **(G)** Representative images of RD cell cultures after treatment with vehicle alone or indicated concentrations of 5Z7O followed by immunostaining for MyHC protein. Nuclei were stained with DAPI. Scale bar: 20 µm. **(H)** Immunoblots presented here demonstrate protein levels of MyHC, myogenin, and GAPDH in RD cells treated with indicated concentrations of 5Z7O.

We also studied the effect of siRNA-mediated silencing of TAK1 on the differentiation of RD cells. Cultured RD cells were transfected with either control or TAK1 siRNAs, and differentiation was assessed 72 h later by immunostaining and immunoblotting for muscle differentiation markers. Like the shRNA approach, siRNA-mediated knockdown of TAK1 enhanced myogenic differentiation in RD cell cultures (**Fig. S3A–C**). To further assess TAK1 function in a system allowing temporal control of gene silencing, we generated lentiviral particles enabling doxycycline-inducible (Tet-On) expression of TAK1 shRNA. This system allows evaluation of differentiation independently of potential proliferation effects. Results showed that doxycycline treatment effectively reduced TAK1 protein levels in RD cultures transduced with Tet-On TAK1 shRNA (**Fig. S3D**) and led to increased expression of myogenin and a higher proportion of MyHC cells (**Fig. S3D-F**).

In parallel, pharmacological inhibition of TAK1 using 5Z7O similarly enhanced myogenic differentiation, evidenced by an increased number of MyHC cells and elevated levels of MyHC and myogenin proteins in RD cultures (**Fig. 5G, H**). Collectively, these results demonstrate that both genetic and pharmacological inhibition of TAK1 enhance myogenic differentiation in RMS cell lines.

### TAK1 inhibits myogenic differentiation in RMS cells through augmenting YAP1 levels

The Hippo-YAP1 signaling pathway plays a significant role in RMS by promoting proliferation and blocking differentiation, particularly in ERMS. YAP1 is a key effector of the Hippo pathway and its hyperactivity is directly linked to ERMS development (24). When the Hippo pathway is active, multiple upstream signals regulate the phosphorylation of MST1/MST2, LATS1/LATS2 kinases, and phosphorylates YAP1 protein which leads to its proteolytic degradation. When the Hippo signaling pathway is inactive, YAP1 is not phosphorylated, leading to its increased levels and translocation to the nucleus where it forms a complex with transcription factor TEADs to augment the gene expression of molecules involved in cell growth and proliferation (51). Intriguingly, RNA-Seq dataset analysis suggested that the Hippo-YAP1 pathway is one of the strongly impacted signaling pathways by knockdown of TAK1 in RD cells (**Fig. 6A**). Furthermore, RPPA analysis showed that the levels of YAP1 were significantly reduced in TAK1 knockdown RD and RH30 cells compared with corresponding control cells (**Fig. 6B**). To further understand the role of TAK1 in the regulation of YAP1, we measured the levels of phosphorylated YAP1 (p-YAP1) and total YAP1 proteins in control and TAK1 knockdown RD and RH30 cells. Results showed that knockdown of TAK1 significantly reduced the levels of p-YAP1 and total YAP1 protein in both RD and RH30 cells compared to corresponding control cells (**Fig. 6C-F**). Similarly, our results showed that siRNA-mediated silencing of TAK1 also drastically reduces the levels of p-YAP1 and YAP1 protein in RD cells (**Fig. 6G, H**)

**FIGURE 6.**
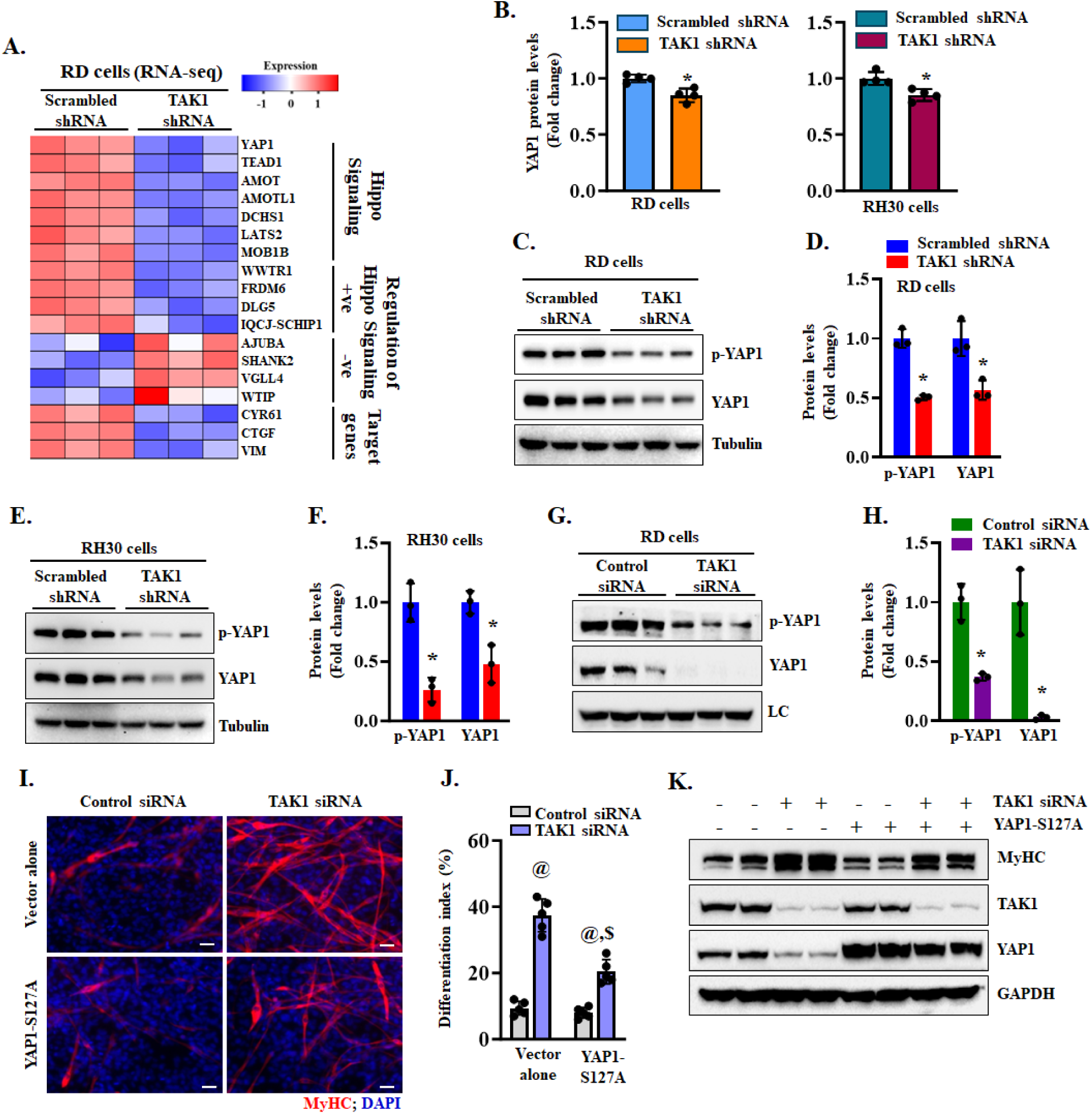
TAK1 inhibits RMS cell differentiation through upregulating YAP1 levels. **(A)** Heatmap generated from RNA-seq dataset showing deregulation of various genes involved in Hippo-YAP1 signaling in TAK1 knockdown RD cultures. **(B)** Analysis of YAP1 protein levels in RPPA dataset of control and TAK1 knockdown RD and RH30 cells. **(C)** Representative immunoblots, and **(D)** densitometry analysis of the levels of p-YAP1 and YAP1 protein in control and TAK1 knockdown RD cells. **(E)** Immunoblots, and **(F)** densitometry analysis of the levels of p-YAP1 and YAP1 protein in control and TAK1 knockdown RH30 cells. n=3 biological replicates in each group. Results are presented as mean ± SD. *p<0.05, values significantly different from corresponding cultures expressing scrambled shRNA by unpaired two-tailed t-test. **(G)** Representative immunoblots, and **(H)** densitometry analysis showing levels of p-YAP1 and YAP1 protein in RD cell cultures transfected with control or TAK1 siRNA. n=3 biological replicates in each group. Data are presented as mean ± SD. *p<0.05, values significantly different from cultures transfected with control siRNA by unpaired two-tailed t-test. **(I)** Representative MyHC-stained images of RD cultures transfected with control or TAK1 siRNA along with vector alone or YAP1-S127A cDNA. Scale bar: 100 µm. **(J)** Quantification of differentiation index in RD cell cultures transfected with control and TAK1 siRNA along with vector alone or YAP1-S127A cDNA. n = 3 biological replicates in each group. Results are presented as mean ± SD. ^@^p<0.05, values significantly different from cultures transfected with control siRNA and vector alone by unpaired two-tailed t-test. ^$^p < 0.05, values significantly different from RD cell cultures transfected with TAK1 siRNA and vector alone by unpaired two-tailed t-test. **(K)** Immunoblots presented here demonstrate the levels of MyHC, TAK1, YAP1, and GAPDH in RD cell cultures transfected with control or TAK1 siRNA with or without YAP1-S127A cDNA. LC, loading control.

To understand the role of the YAP1 downstream of TAK1, we investigated the effect of overexpression of a phosphorylation resistant form of YAP1 (i.e., YAP1-S127A) (52). For this experiment, RD cells were transfected with vector alone or YAP1-S127A cDNA. After 24 h, the cells were transfected with control or TAK1 siRNA oligonucleotides and myogenic differentiation was evaluated 48 h later by immunostaining for MyHC protein. Interestingly, we found that the overexpression of YAP1-S127A protein diminished the effects of knockdown of TAK1 on myogenic differentiation in RD cultures (**Fig. 6I, J**). Western blot analysis confirmed reduced levels of TAK1 protein and increased levels of MyHC in TAK1 siRNA transfected cultures and reduced levels of MyHC and increased YAP1 protein in cultures transfected with TAK1 siRNA along with YAP1-S127A cDNA (**Fig. 6K**). These results suggest that TAK1 inhibits the myogenic differentiation in RMS cells, at least in part, through increasing the levels of YAP1.

### Inducible knockdown of TAK1 inhibits RMS growth in vivo

We next sought to determine whether inhibition of TAK1 attenuates the growth of RMS tumors in vivo. We first generated RD cells with stable expression of Tet-On TAK1 shRNA. To monitor tumor growth, the cells were also transduced with lentiviral particles expressing luciferase cDNA. Finally, the cells were resuspended in Matrigel (BD Biosciences) and injected subcutaneously (5 x 10^6^ cells per mouse) into the flanks of 6-week-old female *Nu/Nu* mice. Mice were monitored twice weekly, and the formation and growth of RD tumors were detected by bioluminescence imaging using a Xenogen IVIS®-system (PerkinElmer, Inc.). When the tumor size reached ∼50 mm^3^, the mice were randomly divided into two groups. One group was fed with normal chow (vehicle) whereas the other group received doxycycline-containing chow. While there was no significant difference in the overall body weight, feeding doxycycline chow significantly reduced RD xenograft growth in mice (**Fig 7A-C**). After 28 days of starting doxycycline diet, the mice were euthanized, and tumor samples were collected for histological and biochemical analysis. Consistent with bioluminescence imaging, wet weight of tumors was significantly reduced in doxycycline-fed mice compared with those fed with normal chow (**Fig. 7D, E**).

**FIGURE 7.**
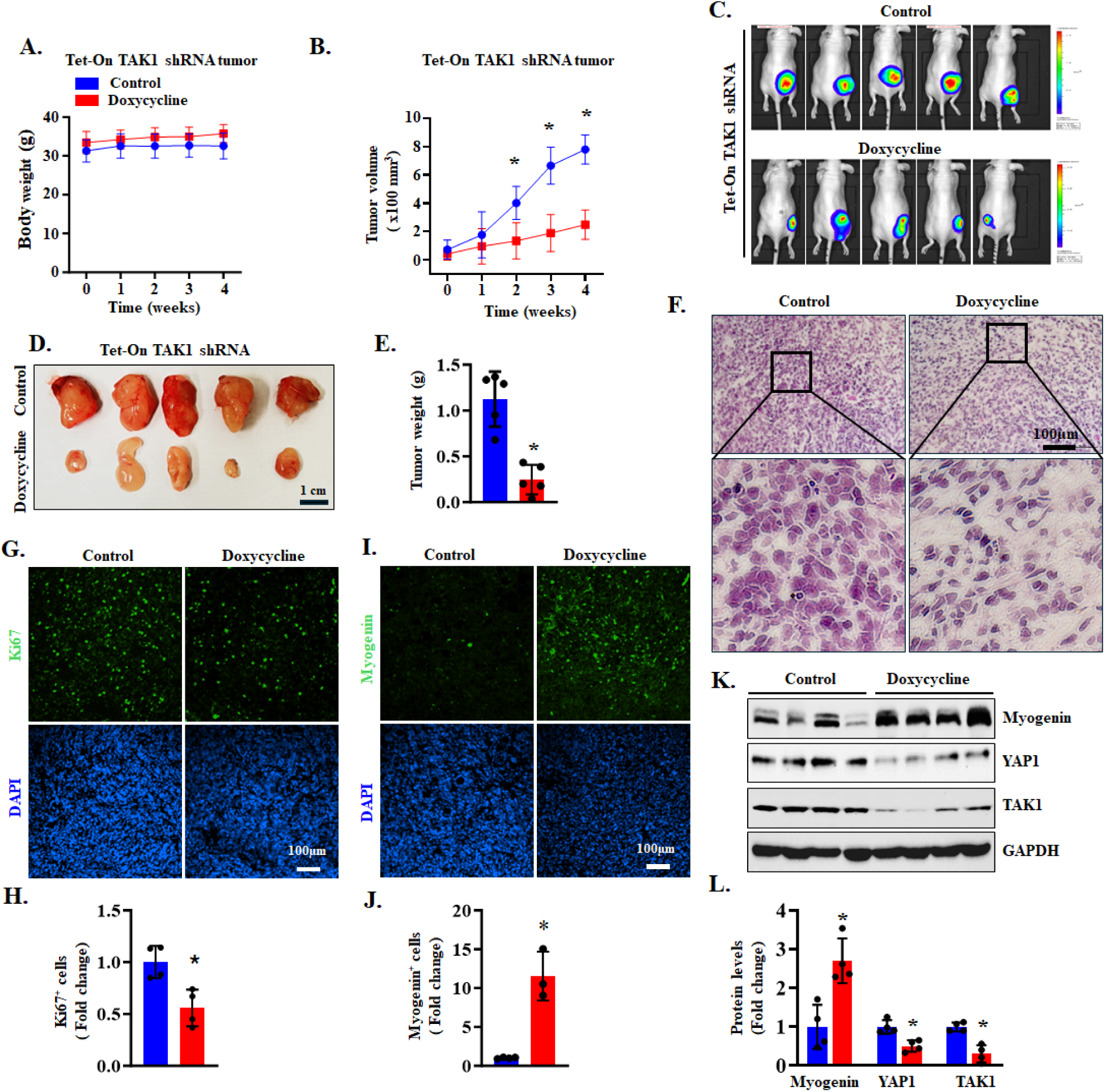
TAK1 knockdown inhibits tumor growth in RD xenograft. **(A)** Body weight of mice inoculated with Tet-On TAK1 shRNA expressing RD cells fed with normal chow or Doxycycline-containing chow. **(B)** Average tumor volume in mice fed with normal chow or Doxycycline containing chow. n= 5 mice in each group. Data are presented as mean ± SD. *p < 0.05, values significantly different from normal chow fed diet at indicated time points by unpaired two-tailed t-test. **(C)** Bioluminescence imaging showing presence RD xenograft in control and doxycycline-treated nude mice. **(D)** Images of tumors at the time of euthanizing the mice between control and doxycycline-treated groups. **(E)** Quantification of wet weight of tumors in control and doxycycline-treated mice. n= 5 mice in each group. Data are presented as mean ± SD. *p < 0.05, values significantly different from control group by unpaired two-tailed t-test. **(F)** Representative images of tumor sections after performing H&E staining. Scale bar: 100µm. **(G)** Representative images of tumor sections after performing immunostaining for Ki67 protein. Nuclei were counterstained with DAPI. Scale bar: 100 µm. **(H)** Quantification of relative number of Ki67^+^ cells in tumor samples of control and doxycycline-treated mice. n= 4 mice in each group. Data are presented as mean ± SD. *p< 0.05, values significantly different from control (normal chow) group by unpaired two-tailed t-test. **(I)** Representative images of tumor sections after performing immunostaining for myogenin protein. Nuclei were counterstained with DAPI.. Scale bar: 100 µm. **(J)** Quantification of relative number of myogenin^+^ cells in tumor samples of control and doxycycline-treated mice. n = 4 for the control group and n = 3 for the doxycycline-treated group. Data are presented as mean ± SD. *p< 0.05, values significantly different from normal chow fed diet at indicated time points by unpaired two-tailed t-test. **(K)** Immunoblots showing relative levels of myogenin, YAP1, TAK1, and GAPDH in tumor samples of control and doxycycline-treated mice. **(L)** Densitometry quantification of levels of myogenin, YAP1 and TAK1 in tumor samples in the two groups. n=4 in each group. Data are presented as mean ± SD. *p<0.05, values significantly different from control group by unpaired two-tailed t-test.

We next generated frozen sections of tumor samples and performed histochemical analysis. H&E staining of tumor sections showed that tumors expressing TAK1 shRNA contained less dense tumor cells and increased abundance of multinucleated cells suggesting growth arrest and myogenic differentiation (**Fig. 7F**). Immunostaining tumor sections for Ki67 protein, a marker for cellular proliferation, showed that there was a significant reduction in the number of Ki67-positive cells suggesting an inhibition in the proliferation of tumor cells following inducible knockdown of TAK1 (**Fig. 7G, H**). Furthermore, there was a significant increase in the number of myogenin-positive cells in TAK1 knockdown tumors compared with controls (**Fig. 7I, J**) suggesting an increase in myogenic differentiation. Our western blot analysis confirmed that the levels of TAK1 protein were reduced in RD tumors of doxycycline-treated mice compared with controls. Furthermore, there was a marked increase in the levels of myogenin and reduction in YAP1 protein in TAK1 knockdown tumor samples compared with controls (**Fig. 7K, L**). Collectively, these results suggest that silencing of TAK1 inhibits proliferation and induces differentiation of RD cells in vivo.

## Discussion

RMS comprises a heterogeneous group of neoplasms marked by uncontrolled cell proliferation and impaired skeletal muscle differentiation. Emerging evidence indicates that RMS is driven by aberrant regulation of various signaling pathways involved in tumorigenesis, proliferation, and myogenic differentiation (8, 10, 19). However, the upstream mechanisms coordinating these pathways remain poorly understood. In this study, we investigated the role of the TAK1 signalosome in regulating RMS growth both in vitro and in vivo. We found that TAK1 is markedly upregulated in both ERMS and ARMS cell lines and in RMS patient tumor samples. Molecular or pharmacological inhibition of TAK1 significantly reduced cell survival, proliferation, and invasiveness, while promoting terminal differentiation in RMS cells. Furthermore, inducible knockdown of TAK1 in xenograft models effectively suppressed tumor growth, likely through reduced proliferation and enhanced differentiation.

Proinflammatory signaling molecules such as TNFα, IL-1β, and agonists of Toll-like receptors (TLRs) activate TAK1, which subsequently triggers downstream signaling cascades, including JNK, p38 MAPK, and the IκB kinase (IKK) complex. This leads to the activation of transcription factors AP-1 and NF-κB, which promote inflammation, cellular proliferation, and migration. TAK1 has been implicated in poor clinical outcomes and tumor metastasis (27). For instance, TAK1 is frequently upregulated in high-grade and metastatic ovarian cancers, where its inhibition suppresses tumor growth via blockade of NF-κB signaling (35). In breast carcinoma, TAK1 inhibition reduces bone metastasis and osteolysis (33), while nanoparticle-mediated delivery of the TAK1 inhibitor 5Z7O significantly diminishes lung metastasis of triple-negative breast cancer (TNBC) in multiple animal models, an effect that is mediated through inhibition of p38 MAPK signaling (34). Similarly, TAK1 inhibition has been shown to reduce proliferation in pancreatic ductal adenocarcinoma cells by targeting the MAPK and NF-κB pathways (53).

Interestingly, the RAS/MEK/ERK, PI3K/AKT, and NF-κB signaling pathways are dysregulated in both ERMS and ARMS cells. Mutations in genes such as RAS, PIK3CA, and PTEN are commonly observed in ERMS, implicating these pathways in RMS pathogenesis (7, 8, 10). Although these signaling cascades are known to support RMS cell proliferation and survival, the upstream mechanisms responsible for their activation and their direct roles in RMS growth and metastasis remain unclear (54–56). Our findings indicate that silencing TAK1 significantly inhibits the proliferation of both ARMS and ERMS cells, suggesting that TAK1 functions as a key upstream regulator of RMS growth. This is further supported by RPPA analysis, which revealed decreased levels of phosphorylated Rb and Ki67, along with upregulation of the cell cycle inhibitor p21, in TAK1 knockdown RMS cells. Moreover, TAK1 knockdown markedly reduced phosphorylation of JNK, the NF-κB subunit p65, and c-Fos (a component of the AP-1 transcription factor) in both RD and RH30 cell lines (**Fig. 2**). These results suggest that TAK1 promotes RMS cell proliferation through activation of the MAPK and NF-κB signaling pathways.

In addition to its role in promoting cellular proliferation, TAK1 also delivers pro-survival signals and inhibits both apoptosis and necroptosis (26). We previously reported that TAK1 is essential for the survival of muscle progenitor cells (39). One mechanism by which TAK1 supports cell survival is through the activation of NF-κB-mediated transcription. In our study, TAK1 knockdown had only a modest effect on RMS cell viability, with 80–90% of cells remaining viable (**Fig. 3**). This contrasts with findings in other cancer types, where NF-κB inhibition leads to a marked reduction in cell viability (57, 58). The reasons for these differential responses are not fully understood. However, a recent study suggests that RMS cells may evade apoptosis through expression of the myogenic transcription factor MyoD, which promotes partial differentiation and represses the expression of the pro-apoptotic gene CYLD (59). This mechanism may allow RMS cells to tolerate reduced NF-κB signaling while maintaining survival.

Despite multimodal therapies, patients with high-risk metastatic RMS continue to face a poor prognosis, with a 5-year overall survival rate of only 30%, a statistic that may reflect the incomplete understanding of the mechanisms driving RMS metastasis (4, 6, 9). The EMT is a key process in tumor metastasis across various cancer types, including RMS. EMT is characterized by the loss of E-cadherin expression or function, a significant reduction in tight junction proteins, and an upregulation of mesenchymal markers such as N-cadherin (60). The initiation of EMT is primarily driven by the activation of specific transcription factors, including the zinc-finger proteins Snail1 and Slug, zinc finger E-box-binding homeobox 1 (ZEB1), ZEB2, and Twist. These transcription factors repress the gene expression of various molecules involved in cell-cell adhesion by binding to their promoter regions (50, 61). Previous animal studies have shown that TAK1 plays an important role in metastasis of breast carcinoma cells and TNBC (33, 34). Similarly, our experiments in the present study demonstrate that the silencing of TAK1 also inhibits the migration and invasiveness of both ARMS and ERMS cell lines. Although exact mechanisms remain unknown, our results demonstrate that knockdown of TAK1 in RD or RH30 cells reduces the levels of key EMT-related transcription factors (e.g., Twist, Zeb1, Smad2), signaling molecules, and cell surface molecules involved in cell migration and metastasis suggesting that TAK1 could be a potential target to prevent RMS metastasis (**Fig. 4**).

Deregulation of myogenic differentiation is a key driver of the uncontrolled proliferation observed in RMS tumor cells (5, 8). However, the molecular and signaling mechanisms underlying this impaired differentiation remain poorly understood and appear to be highly heterogeneous likely reflecting the diversity of oncogenic drivers associated with distinct RMS subtypes. MyoD, a master transcription factor, plays a central role in committing muscle progenitor cells to the myogenic lineage (62, 63). In RMS, MyoD transactivation is frequently impaired, possibly due to interactions with transcriptional repressors or competition for E-box binding sites in the promoters of myogenic genes (8, 64–67). In addition to transcriptional repression, increasing evidence suggests that altered signaling pathways also contribute to defective myogenesis in RMS (8, 10, 19). For example, a recent study demonstrated that MAPK ERK2 suppresses myogenic differentiation in ERMS by downregulating the expression of MYOG. Treatment with trametinib, a MEK inhibitor, was shown to displace ERK2 from the MYOG promoter, thereby relieving transcriptional repression and promoting terminal differentiation (68). In our study, we found that TAK1 inhibition enhances differentiation in both ERMS and ARMS cell lines. Notably, TAK1 knockdown also reduced phosphorylation of ERK1/2 in RH30 cells (**Fig. S2**), suggesting that one potential mechanism by which TAK1 impairs myogenic differentiation is through activation of the MAPK signaling pathway.

YAP/TAZ proteins are frequently hyperactivated in cancer due to dysregulation of the Hippo signaling pathway. This leads to their nuclear accumulation, where they drive the expression of genes that promote proliferation, inhibit apoptosis, and support tumor progression (51). In RMS, YAP1 is upregulated in both ERMS and ARMS tumors, where it contributes to tumorigenesis by promoting proliferation and suppressing myogenic differentiation and apoptosis. In vivo suppression of YAP1, either through RNA interference or pharmacological inhibition with the YAP-TEAD inhibitor verteporfin, has been shown to reduce RMS tumor growth (24, 69). Our findings suggest that one mechanism by which TAK1 inhibits myogenic differentiation is through stabilization of YAP1 protein. Specifically, TAK1 knockdown reduced both phosphorylated and total YAP1 protein levels in ERMS and ARMS cell lines. Furthermore, overexpression of a degradation-resistant YAP1 mutant partially reversed the differentiation-promoting effects of TAK1 inhibition in RD cells (**Fig. 6**), supporting a functional link between TAK1 and YAP1 stability in RMS. Although the precise mechanism by which TAK1 regulates YAP1 levels in RMS remains unclear, previous studies have shown that phosphorylated TAK1 binds and stabilizes YAP1/TAZ in bone marrow-derived mesenchymal stem cells (70). In pancreatic cancer cells, TAK1 enhances YAP1/TAZ stability by promoting K63-linked ubiquitination while inhibiting K48-linked ubiquitination, which typically targets proteins for proteasomal degradation (71). It is therefore plausible that TAK1 similarly stabilizes YAP1 in RMS cells through modulation of ubiquitin signaling.

Our in vivo results further support the critical role of TAK1 in sustaining RMS tumor growth and maintaining the undifferentiated, proliferative state of RMS cells (**Fig. 7**). Using a doxycycline-inducible shRNA system in RD xenografts, we demonstrate that inducible TAK1 knockdown significantly attenuates tumor growth without affecting overall body weight, indicating that the observed effects are not due to systemic toxicity. The role of TAK1 in RMS growth is also supported by our histological and immunohistochemical analyses which revealed profound changes in tumor architecture and cellular behavior following TAK1 knockdown. H&E staining showed reduced cellular density and an increase in multinucleated cells which are hallmarks of myogenic differentiation and growth arrest (**Fig. 7**). This was further supported by a significant reduction in Ki67^+^ cells, indicative of reduced cellular proliferation, and a corresponding increase in myogenin^+^ cells, suggesting enhanced commitment to the myogenic lineage. Notably, TAK1 suppression led to an upregulation of myogenin, a key transcription factor required for terminal muscle differentiation, along with a reduction in YAP1 protein levels (**Fig. 7**). Since YAP1 is known to repress myogenic differentiation and promote RMS proliferation (24, 69), these results suggest that TAK1 maintains the undifferentiated state of RMS cells in part through stabilization of YAP1. This is consistent with our in vitro findings and previously published studies also implicating TAK1 in YAP1 regulation through post-translational mechanisms (70, 71).

In summary, this study provides initial evidence that TAK1 is not only essential for RMS cell proliferation but also actively suppresses myogenic differentiation. Targeting TAK1 may therefore offer a dual therapeutic benefit by simultaneously inhibiting tumor growth and promoting differentiation of RMS cells, a strategy that could complement current treatments focused on eliminating proliferative tumor cells. Furthermore, differentiation-based therapies for RMS may offer significant advantages over conventional chemotherapy by reducing adverse side effects. Future studies will assess the effects of small-molecule TAK1 inhibitors on the growth and metastasis of orthotopic patient-derived RMS xenografts. In parallel, it will be important to elucidate the precise molecular mechanisms through which TAK1 regulates YAP1 stability and to determine whether combined targeting of TAK1 and other oncogenic pathways can further enhance therapeutic efficacy in RMS.

## Materials and Methods

### Cell lines

RD, RH30, and HTB82 cells lines were purchased from American Type Culture Collection (ATCC, Manassas, Virginia). RH36 and RH41 cells were kindly provided by Dr. Peter J Houghton of The University of Texas Health Science Center at San Antonio, Texas. The RD and RH36 cells were maintained in DMEM, The RH30 and RH41 cells were maintained in RPMI and HTB82 cell was maintained in McCoy’s 5A medium, all supplemented with 10% fetal bovine serum and 1x penicillin-streptomycin. The absence of mycoplasma contamination was confirmed using the Universal Mycoplasma Detection Kit (ATCC, Cat. No. 30-1012K). Control and TAK1 siRNA were purchased from Santa Cruz Biotechnology, Inc. (Dallas, TX). The cells were transfected using Lipofectamine® RNAiMAX reagent (Invitrogen).

### Generation of lentiviral particles

Lentiviral particles expressing scrambled or TAK1 shRNA were generated following a protocol as previously described (72). In brief, the target siRNA sequence for human TAK1 mRNA was identified using BLOCK-iT™ RNAi Designer online software (Life Technologies). The shRNA oligonucleotides were synthesized to contain the sense strand of target sequences for human TAK1 (i.e. shRNA seq 1: GCT GAA CCA TTG CCA TAT TAT or shRNA Seq 2: GCA ACC CAA AGC GCT AAT TCA), short spacer (CTCGAG), and the reverse complement sequences followed by five thymidines as an RNA polymerase III transcriptional stop signal. Oligonucleotides were annealed and cloned into pLKO.1-mCherry-Puro with AgeI/EcoRI sites. For generation of Tet-ON TAK1 shRNA, the TAK1 shRNA seq 1 oligos were cloned in Tet-pLKO-puro plasmid (Addgene, Plasmid #21915). The insertion of shRNA in the plasmid was confirmed by DNA sequencing. For generation of lentiviral particles, the HEK293T cells were co-transfected with 5 μg psPAX2 (Addgene, Plasmid # 12260), 5 μg pMD2.G (Addgene, Plasmid # 12259) and 10 μg of pLKO.1-mCherry-scrambled shRNA or pLKO.1-mCherry-TAK1 shRNA using PEI reagent (Sigma-Aldrich, USA). After 8 h of transfection, the media was replaced with fresh media. Lentiviral particles were collected 48 h after transfection and filtered through 0.45-micron filters. Cultured RMS cells were transduced with lentiviral particles containing scrambled or TAK1 shRNA in growth medium containing 6-8 μg/ml polybrene for 24 h. For generation of stable Tet-on Scrambled shRNA or TAK1 shRNA expressing cells, the transduced cells were grown for 72 h and selected in the presence of 1µg/ml puromycin.

### Cell proliferation assay

The effect of TAK1 knockdown on RMS cell proliferation was evaluated as previously described (73). Briefly, cells were seeded at 1,000 cells per well in a PhenoPlate™ 96-well tissue culture plate (Revvity, USA). The plate was scanned using the EnSight Multimode Plate Reader equipped with well-imaging technology (PerkinElmer, MA, United States). Cell count was obtained by digital phase and bright field imaging.

### EdU incorporation and fluorescence-activated cell sorting (FACS) analysis

The cell proliferation was also examined using an EdU incorporation assay with the Click-iT™ EdU Cell Proliferation Kit for Imaging, Alexa Fluor™ 488 dye (Invitrogen, USA). Briefly, RMS cells were transduced with either scrambled shRNA or TAK1 shRNA lentivirus. The following day, the medium was replaced with fresh culture medium, and the cells were cultured for an additional three days. Subsequently, 1 µM EdU was added, and the cells were incubated for 2–3 hours. After EdU incubation, the cells were collected using trypsin and processed according to the manufacturer’s instructions for the EdU detection kit. EdU^+^ cells were analyzed by flow cytometry using a C6 Accuri cytometer (BD Biosciences), and the data were processed using FlowJo software (version 10.10.0, RRID:SCR_008520).

### MTT assay

Cell growth and viability of control and TAK1 knockdown RMS cells were assessed using the MTT assay. Briefly, 5 × 10³ cells per well were seeded in 96-well plates and cultured for 5 days. The cells were then incubated at 37°C for 4 hours with MTT solution at a final concentration of 0.5 mg/ml. After incubation, the culture medium was removed, and 100 µl of DMSO was added to each well to solubilize the formazan crystals. The optical density was measured at 570 nm using a microplate reader (SpectraMax® i3x). The results were expressed as the relative viability of TAK1 knockdown cells compared to their corresponding controls.

### Clonogenic assay

Anchorage-dependent growth of RMS cells was assessed using a colony formation assay. Control and TAK1 knockdown RMS cells (5 × 10³ cells per well) were seeded in 6-well plates and cultured for 2 weeks, with medium replaced every 3 days. Colonies were fixed with pre-chilled methanol/acetone (1:1) at −20°C for 15 min, stained with 0.1% crystal violet for 15 min at room temperature, and rinsed with tap water. For quantification, the bound dye was solubilized in 1 ml of 10% acetic acid per well, shaken for 15 min at room temperature, diluted 1:4 with water, and the absorbance was measured at 590 nm using a SpectraMax® i3x microplate reader.

### Transient transfection and reporter gene activity

Skeletal α-actin promoter reporter gene activity was measured as previously described (40). In brief, control and TAK1 knockdown RD or RH30 cells were plated in 12-well tissue culture plates. The cells were transfected with pGL4 (Promega) or pSK-Luc plasmid using Lipofectamine 2000 transfection reagent according to the protocol suggested by the manufacturer (Invitrogen). Transfection efficiency was controlled by co-transfection of the cells with Renilla luciferase encoding plasmid pRL-TK (Promega). After 3 days, the cells were lysed, and the lysates were processed for luciferase expression using a Dual luciferase assay system per the manufacturer’s instructions (Promega). Luciferase measurements were made using a luminometer (SpectraMax® i3x).

### Annexin V staining and FACS analysis

For analysis of cell viability, we performed Annexin V and propidium iodide (PI) staining followed by FACS analysis using the Annexin V-FITC Apoptosis Staining/Detection Kit (ab14085, Abcam, USA) following a protocol suggested by the manufacturer.

### Transwell migration and invasion assay

For the Transwell migration assay, 5 × 10^4^ control or TAK1 knockdown RD or RH30 cells in 200 μL serum-free DMEM were seeded into the top insert of a Boyden chamber (Corning Inc., Corning, NY, USA), while 800 μL medium with 10% FBS was loaded into the well below. After 24 h of incubation, migrating cells that passed through the filter were stained with 0.1% crystal violet solution. For the Transwell invasion assay, all procedures were similar except that transwell membrane was coated with 100 μl of 300 μg/mL Matrigel (BD Bioscience, San Jose, CA, USA). Finally, the images of the crystal violet-stained cells that passed the filter were captured and the crystal violet signal intensity divided by total area was quantified using the ImageJ software (NIH).

### Immunofluorescence

RMS cells were transduced with lentivirus encoding either scrambled shRNA or TAK1 shRNA and cultured for 5 days post-transduction. After 5 days, cells were washed with PBS, fixed with 4% paraformaldehyde (PFA) for 15 min at room temperature, and permeabilized with 0.5% Triton X-100 in PBS for 15 min. Cells were then blocked with 5% goat serum in PBS containing 0.1% Triton X-100. The anti-MyHC (clone MF20) primary antibody was diluted 1:50 in 5% goat serum and incubated with the cells overnight at 4°C. After washing with PBST, cells were incubated with goat anti-mouse secondary antibody (diluted 1:300 in 5% goat serum) for 1 h at room temperature. Nuclei were counterstained with DAPI for 15 min. Images were acquired using a fluorescence inverted microscope (Nikon Eclipse TE2000-U) equipped with a digital camera (Digital Sight DS-Fi1). The percentage of MyHC cells was quantified using ImageJ software.

### Differentiation index

Differentiation of RMS cells was quantified by measuring the differentiation index which is defined as: (Number of nuclei in MyHC^+^ cells/Total number of nuclei) × 100.

### Western Blot

Cultured RMS cell lines or RD tumor samples were washed with PBS and homogenized in lysis buffer A (50 mM Tris-Cl (pH 8.0), 200 mM NaCl, 50 mM NaF, 1 mM dithiothreitol, 1 mM sodium orthovanadate, 0.3% IGEPAL, and protease inhibitors). Approximately, 20-40 μg protein was resolved on each lane on 8-10% SDS-PAGE gel, transferred onto a PVDF membrane, and probed using a specific primary antibody described in supplemental **Table S1**. Bound antibodies were detected by secondary antibodies conjugated to horseradish peroxidase (Cell Signaling Technology). Signal detection was performed by an enhanced chemiluminescence detection reagent (Bio-Rad). Approximate molecular masses were determined by comparison with the migration of prestained protein standards (Bio-Rad). Uncropped gel images are presented in **Fig. S4**.

### RNA-sequencing and data analyses

Total RNA from control and TAK1 knockdown RD cells was extracted using TRIzol reagent (Thermo Fisher Scientific) using RNeasy Mini Kit (QIAGEN) according to the manufacturers’ protocols. The mRNA-Seq library was prepared using poly (A)-tailed enriched mRNA at Novogene (Sacramento, CA). The Illumina NovaSeq 6000 was used to produce 75 base paired-end mRNA-Seq data at an average read depth of ∼38 M reads/sample. Original image data file from high-throughput sequencing platforms (like Illumina) was transformed into sequenced reads (called Raw Data/Raw Reads) by CASAVA base recognition (Base Calling). To obtain clean reads, the sequencing raw reads were filtered to remove adaptor contamination, constitution of uncertain nucleotides (N>10%), and reads containing more than 50% of low-quality nucleotides (base quality<5). STAR software was used to accomplish mapping and alignment, which precisely and effectively performs positioning junction reads for RNA sequencing data analysis. Illumina sequencing adapters were trimmed, and reads were aligned to the human reference genome Homo Sapiens (GRCh38/hg38). Normalization of RNA-Seq data was performed using the trimmed mean of M-values. Genes with |log_2_FC| ≥ 0.25) and adjusted p-value <0.05 were assigned as differentially expressed genes and represented in a volcano plot using the ggplot function in R software (v 4.2.2). Pathway enrichment analysis associated with the up-regulated and down-regulated genes were identified using the Metascape gene annotation and analysis tool (metascape.org) as described (74, 75). Heatmaps were generated using the heatmap.2 function (76) using z-scores calculated based on transcripts per million (FPKM) values. TPM values were converted to log (FPKM+1) to handle zero values. Genes involved in specific pathways were manually selected for heatmap expression plots.

### Reverse Phase Protein Array (RPPA)

RPPA assays were carried out as described (77). In brief, control and TAK1 knockdown RD or RH30 cells were washed with PBS and lysed in lysis buffer A. Total protein content was quantified, and lysates were submitted to the Antibody-based Proteomics Core at Baylor College of Medicine for RPPA analysis. Each sample was run in technical triplicate, with four biological replicates per experimental group. Fluorescence-labeled slides were scanned using the Molecular Devices GenePix 4400 AL scanner, and signal intensities were extracted using GenePix Pro 7.0. Spot intensities were calculated by subtracting local background signals, followed by group-based normalization to adjust for total protein variation, background noise, and non-specific labeling. For each sample, the median value of the technical triplicates was used for statistical analysis. Proteins with maximum signal intensities below 200 were excluded from further analysis. Differential protein expressions between experimental conditions were assessed using Student’s *t*-test, with a significance threshold of p < 0.05. Heatmaps for visualizing differential expression patterns were generated by R software (v4.4.1) using heatmap.2 function. Significantly differentiated proteins were further mapped to their corresponding gene symbols. The enricher function from the clusterProfiler R package was used for enrichment analysis, with Hallmark gene sets provided as a custom TERM2GENE mapping. Pathways with an adjusted p value < 0.05 were considered significantly enriched.

### Xenograft growth mouse model

Six-week-old male Nu/Nu mice were purchased from Charles River Laboratory. RD cells expressing Tet-On TAK1 shRNA along with luciferase were suspended in Matrigel (Corning, USA), and 5 × 10 cells were injected subcutaneously into the flank of each mouse. Tumor growth was monitored, and tumor length (L) and width (W) were measured weekly. Tumor volume was calculated using the formula: 0.5 × L × W². Tumor growth was also assessed by bioluminescence imaging using a Xenogen IVIS®-system (PerkinElmer, Inc.). When tumor volumes reached approximately 50-100 mm³, mice were randomly divided into two groups. One group was fed normal chow (control), while the other group received doxycycline-containing chow. Mice were euthanized after 4 weeks of doxycycline treatment. Tumors were isolated and processed for histological and biochemical analyses. Tumor samples were fixed in 4% paraformaldehyde overnight at 4°C, washed with cold PBS, and embedded in Frozen Section Compound (FSC22, Leica, USA). Cryosections of 10 µm thickness were prepared and stained with hematoxylin and eosin (H&E). Sections were also immunostained for Ki67 and myogenin. The animal protocol was approved by the Institutional Animal Care and Use Committee (IACUC) of the University of Houston.

### Statistical analysis

Results are expressed as means ± SD. An unpaired, 2-tailed Student’s *t* test was used to compare quantitative data populations with normal distribution and equal variance. The value of *p* < 0.05 was considered significant.

## Data availability

All other raw data generated for this study has been included with this manuscript.

## Authors’ contributions

AK, MVT, BG, and BAK designed the work. ATV, ASJ, AR, KM, PTH, RB, MTS, and TS performed the experiments and analyzed the results. ATV, ASJ, and AK wrote the manuscript. All authors edited and finalized the manuscript.

## Acknowledgements

We thank Dr. Peter J Houghton of The University of Texas Health Science Center at San Antonio, Texas for providing RH36 and RH41 cell lines.

