## Supplementary material for "TAK1 is a key regulator of oncogenic signaling and differentiation blockade in rhabdomyosarcoma": Figures S1-S4 and Table S1

This file contains **Figures S1-S4** and **Table S1**.

### Supplemental Figures and Legends

FIGURE S1

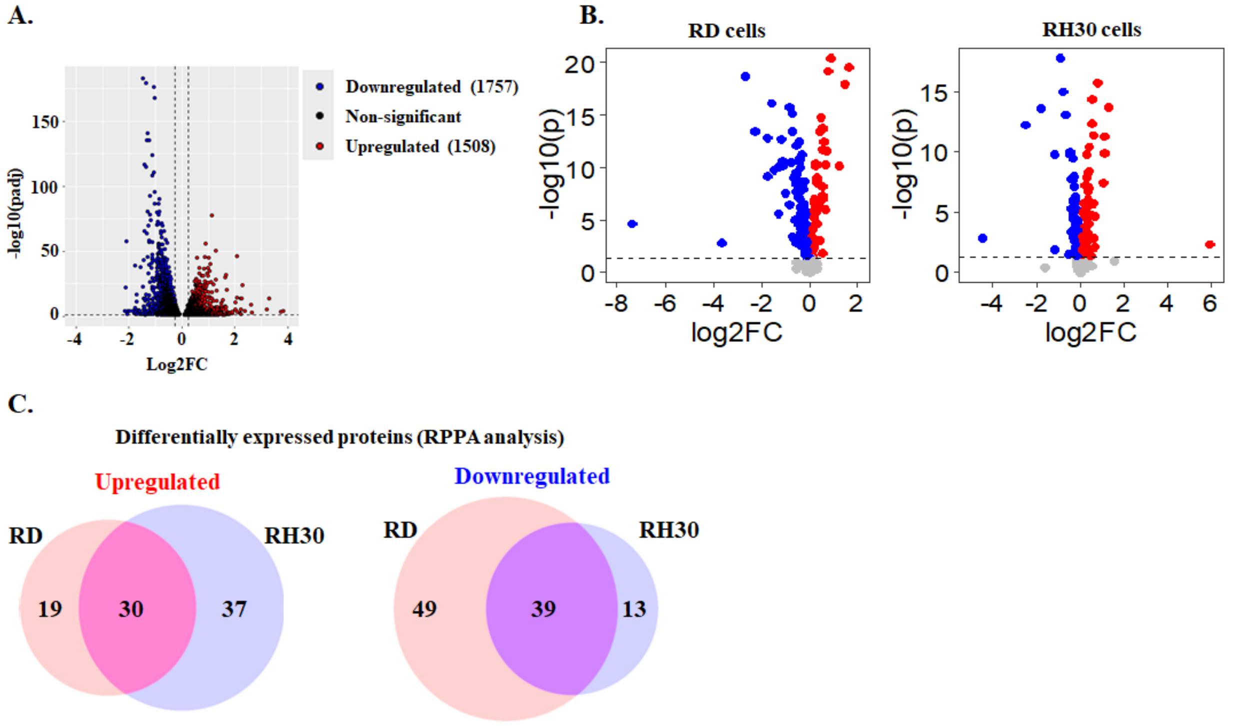


D.

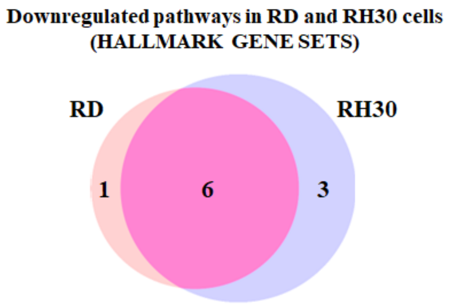


| Common downregulated pathways in TAK1 knockdown RD and RH30 cells |
|---|
| HALLMARK_E2F_TARGETS |
| HALLMARK_PI3K_AKT_MTOR_SIGNALING |
| HALLMARK_EPITHELIAL_MESENCHYMAL_TRANSITION |
| HALLMARK_G2M_CHECKPOINT |
| HALLMARK_GLYCOLYSIS |
| HALLMARK_WNT_BETA_CATENIN_SIGNALING |

**FIGURE S1. Analyis of RNA-Seq and RPPA dataset. (A)** Volcano plot from RNA-seq analysis of TAK1 knockdown versus control RD cells illustrating down-regulated (blue dots) and up-regulated (red dots) genes with a threshold of $\log_2 FC \geq |0.5|$ and $P$-value $\leq 0.05$. (B) Volcano plots of RPPA analysis of TAK1 knockdown RD and RH30 cells versus corresponding control showing down-regulated (blue dots) and up-regulated (red dots) proteins with a threshold of $\log_2 FC > 0$ and $\log_2 FC < 0$ with $P$-value $< 0.05$. **(C)** Vein diagram showing distinct and commonly upregulated and downregulated proteins in TAK1 knockdown RD and RH30 cells. **(D)** Pathway enrichment analysis using Hallmark gene sets in RPPA dataset show common downregulated pathways in TAK1 knockdown RD and RH30 cells.

### FIGURE S2

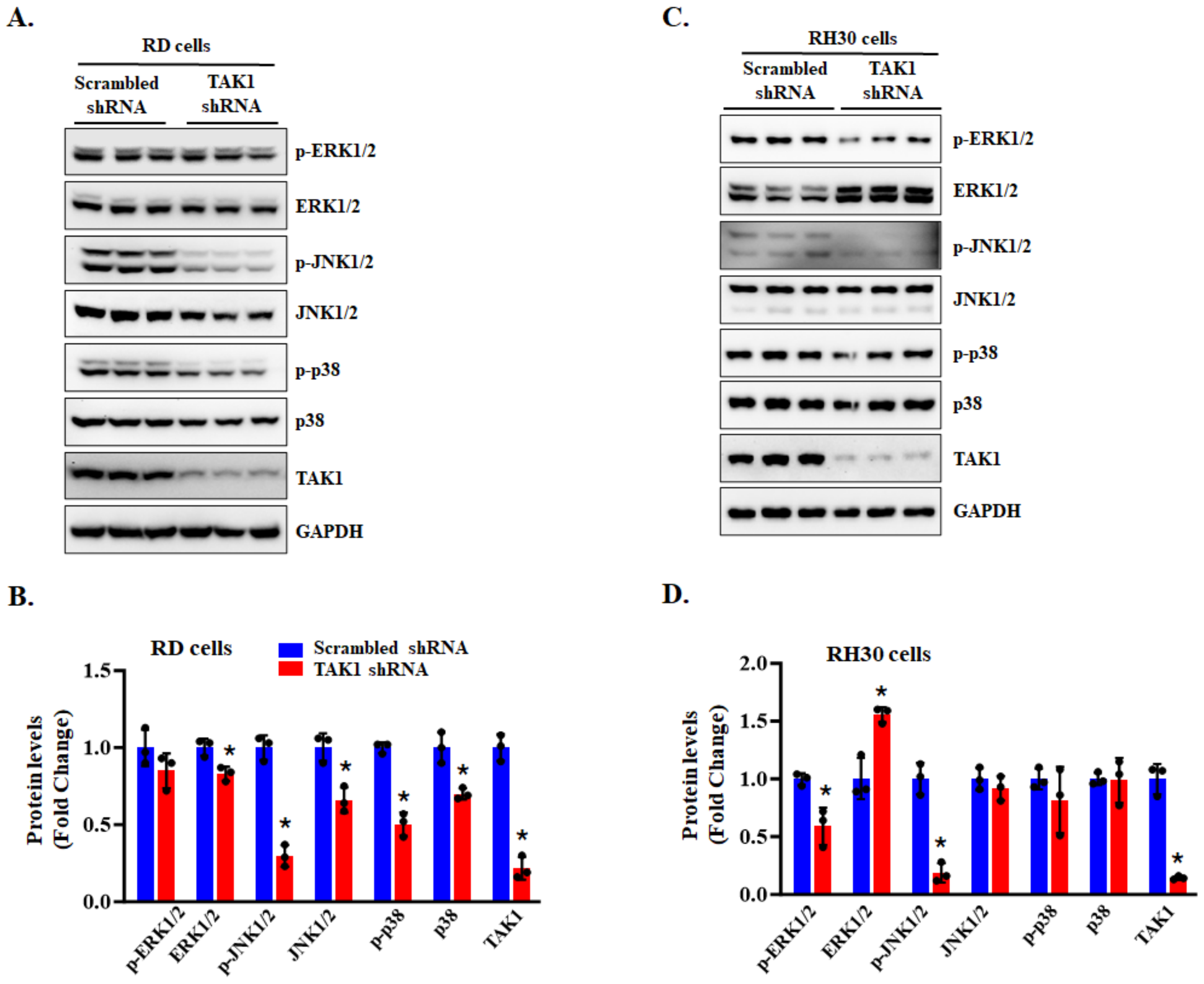


**FIGURE S2. Effect of TAK1 knockdown on the phosphorylation of MAPKs in RMS cells.** (**A**) Immunoblots and **(B)** densitometry analysis demonstrating the levels of phosphorylated and total ERK1/2, JNK1/2, and p38 protein and total TAK1 protein in control and TAK1 knockdown RD cells. (**C**) Immunoblots, and **(D)** densitometry analysis demonstrating the levels of phosphorylated and total ERK1/2, JNK1/2, and p38 protein and total TAK1 protein in control and TAK1 knockdown RH30 cells. n = 3 biological replicates in each group. Data are presented as mean ± SD. *p<0.05 from corresponding cultures expressing scrambled shRNA by unpaired two-tailed t-test.

**FIGURE S3**

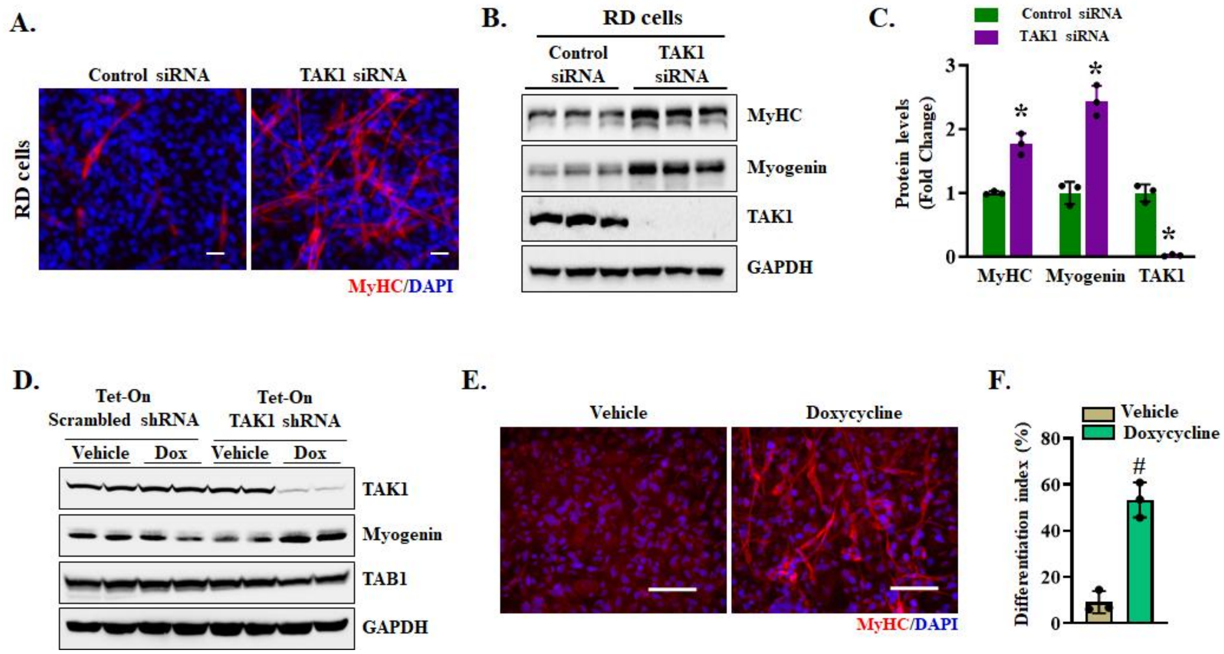


**FIGURE S3. Effect of inhibition of TAK1 in myogenic differentiation of RD cells. (A)** Representative photographs of control and TAK1 siRNA transfected RD cultures after immunostaining for MyHC protein. Nuclei were counterstained with DAPI. Scale bar: 20 µm. **(B)** Immunoblots and **(C)** densitometry analysis demonstrating the levels of MyHC, myogenin, TAK1, and GAPDH protein in RD cell cultures transfected with control or TAK1 siRNA. n=3 biological replicates in each group. Results are presented as mean ± SD. *p<0.05, values significantly different from corresponding cultures transfected with control siRNA. **(D)** Immunoblots presented here demonstrate the levels of TAK1, myogenin, TAB1, and GAPDH in RD cell cultures transduced with lentiviral particles expressing Tet-On scrambled shRNA or Tet-On TAK1 shRNA after treatment with vehicle alone or doxycycline. **(E)** Representative images of RD cell cultures expressing Tet-On TAK1 shRNA after treatment with vehicle alone or doxycycline followed by immunostaining for MyHC protein. Nuclei were stained with DAPI. Scale bar: 100 µm. **(F)** Quantification of differentiation index in control and inducible TAK1 knockdown RD cultures. n = 3 biological replicates in each group. Data are presented as mean ± SD. $^{\#}$p<0.05, values significantly different from cultures treated with vehicle alone by unpaired two-tailed t-test.

**FIGURE S4**

**Figure 1C.**

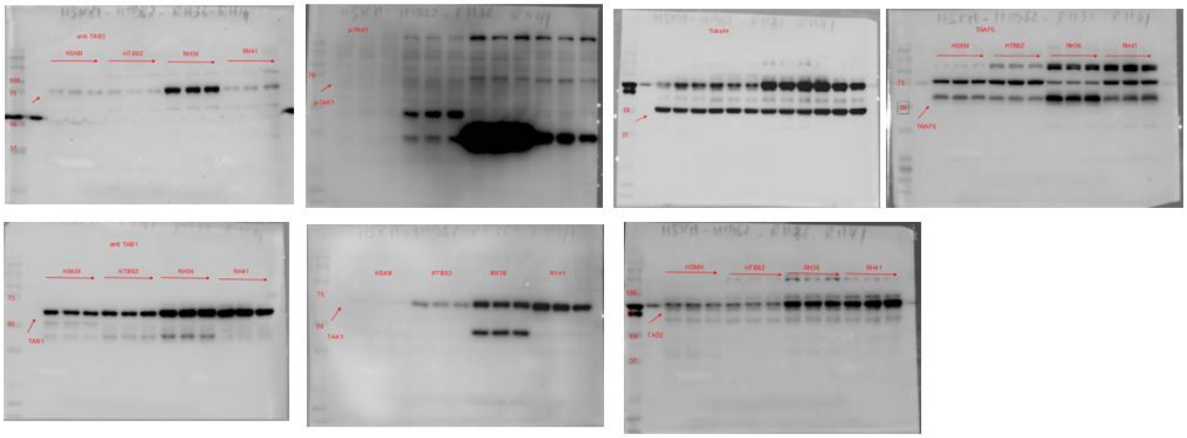

**Figure 1E.**

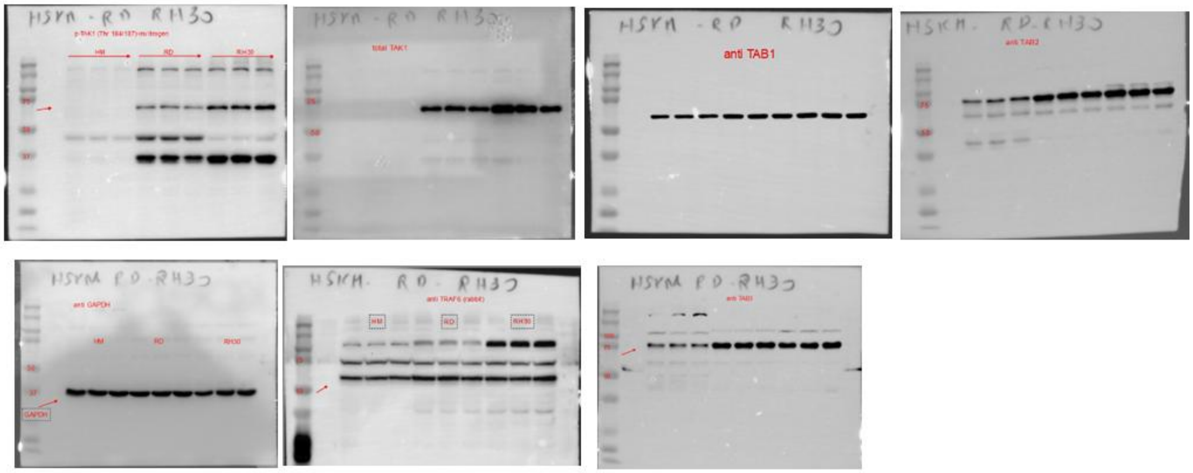


**Figure 2A.**

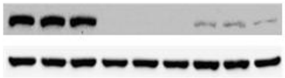

**FIGURE S4** (Continuation)

**Figure 5D.**

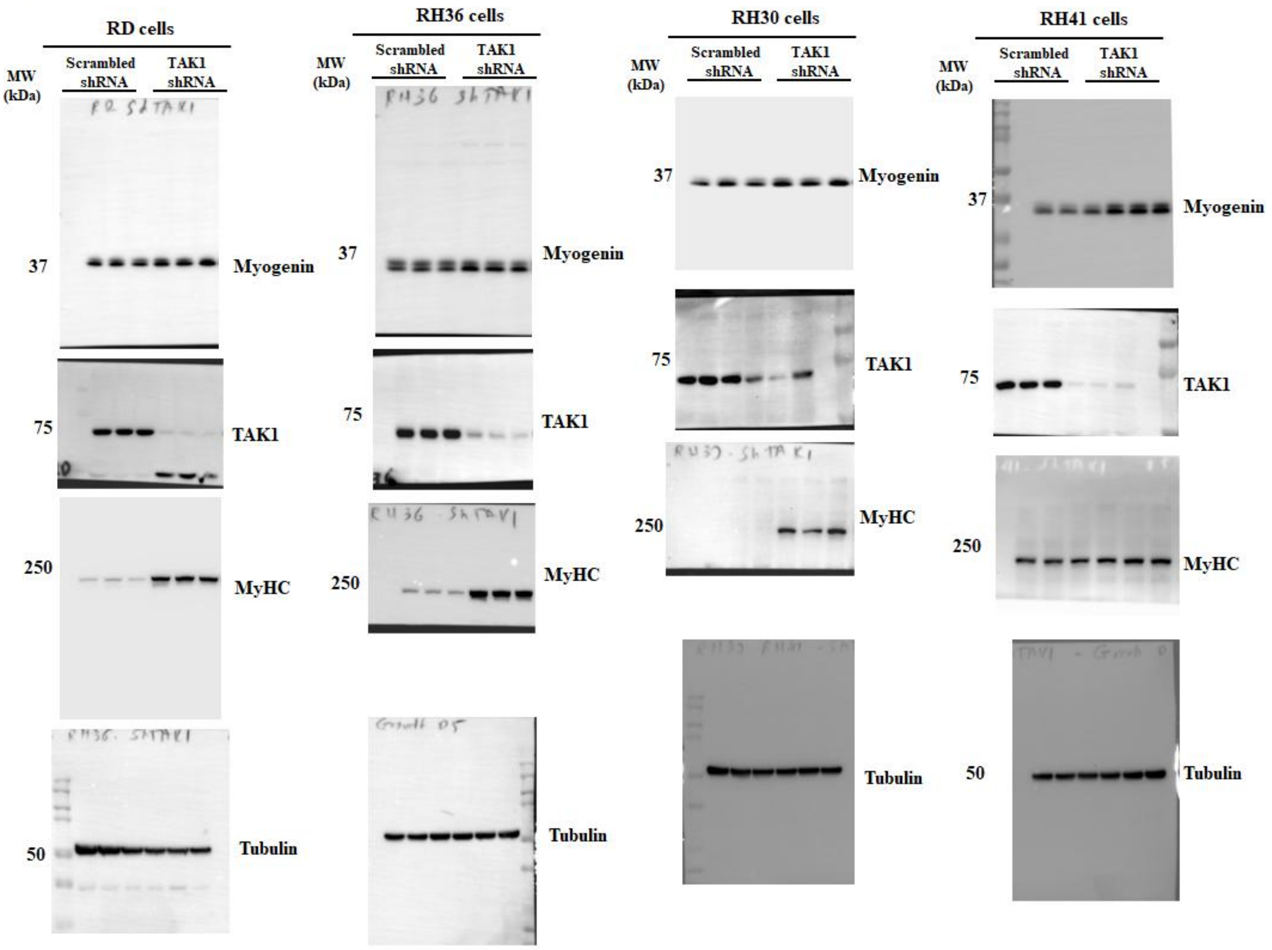


**Figure 5H.**

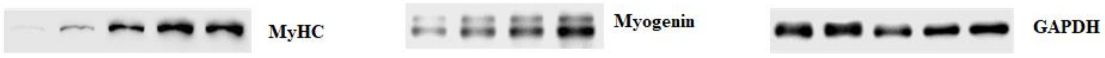

**FIGURE S4** (Continuation)

**Figure 6C.**

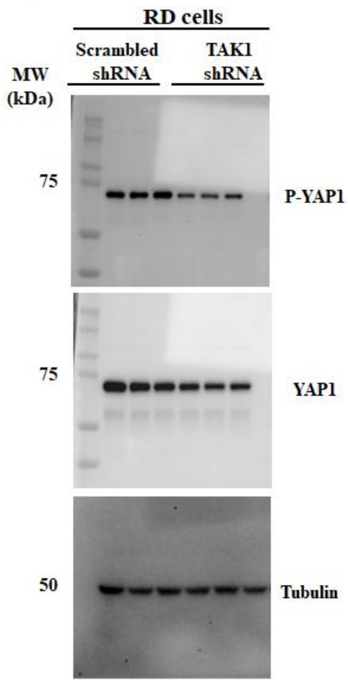


**Figure 6E.**

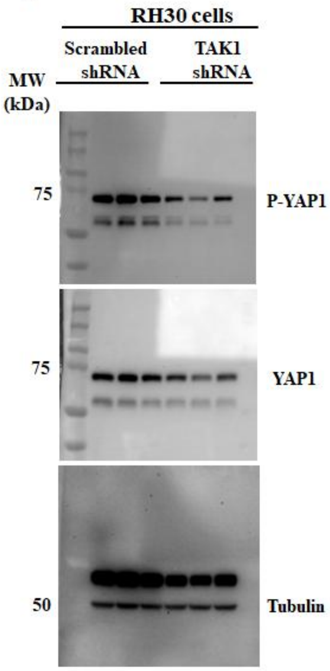


**Figure 6G.**

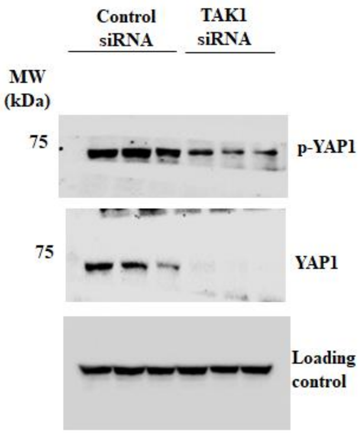


**Figure 6K.**

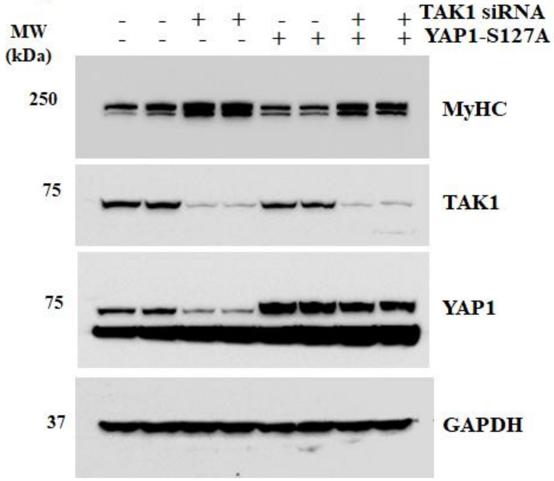


**Figure 7K.**

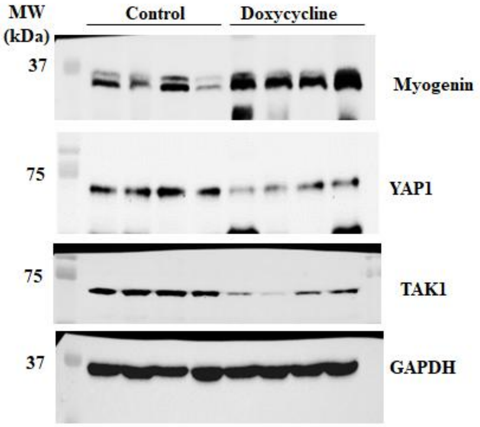

**FIGURE S4** (Continuation)

**Figure S2A.**

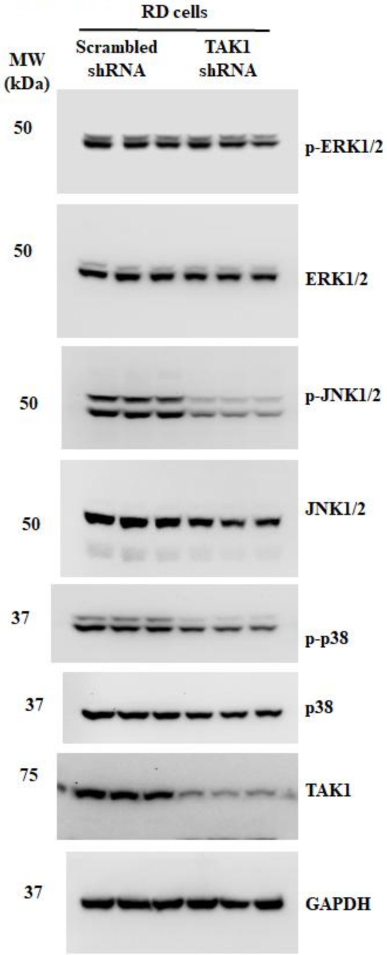


**Figure S2C.**

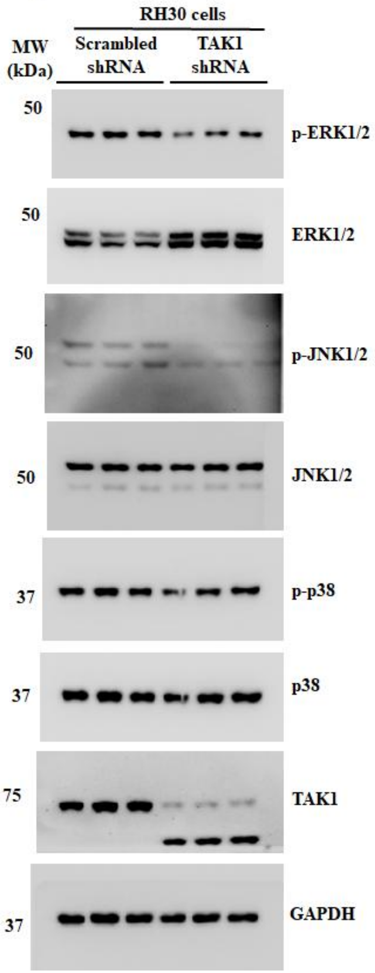


**Figure S3B.**

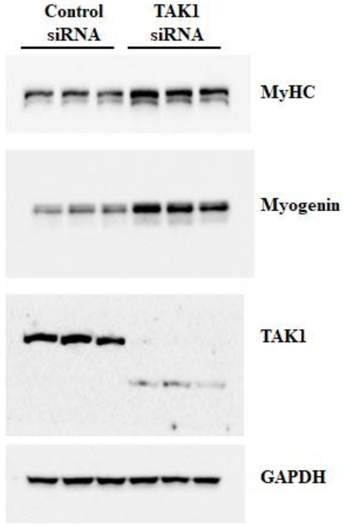

**FIGURE S4** (Continuation)

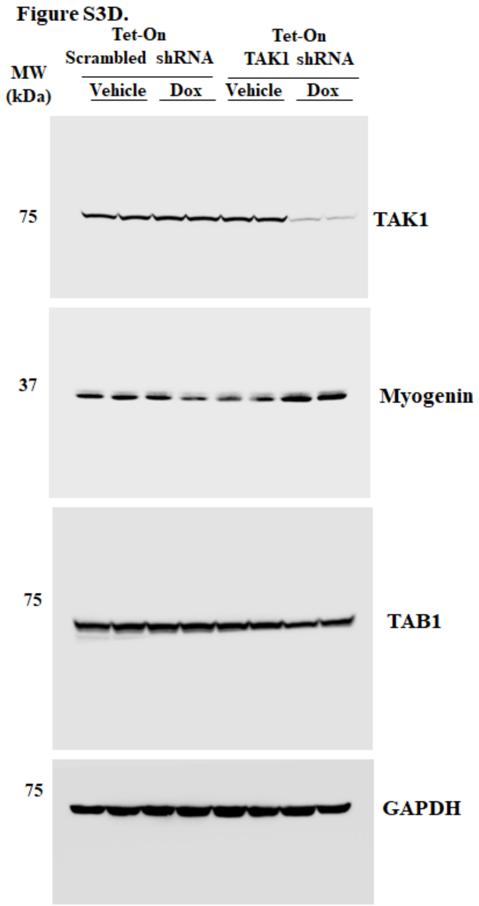


**FIGURE S4. Uncropped western blot images.** Uncropped immunoblot images used in the main figures and supplemental results.

**Table S1. The list of antibodies used in various experiments.**

| Antibody | Dilution | Source | Identifier |
|---|---|---|---|
| Rabbit-anti-p-TAK1 (Thr184/187) | 1:1000 (WB) | Invitrogen | # MA5-15073 |
| Rabbit-anti-TAK1 | 1:1000 (WB) | Cell Signaling Technology | # 5206 |
| Rabbit-anti-TAB1 | 1:1000 (WB) | Cell Signaling Technology | # 3226 |
| Rabbit-anti-TAB2 | 1:1000 (WB) | Cell Signaling Technology | # 3745 |
| Rabbit-anti-TAB3 | 1:1000 (WB) | Cell Signaling Technology | # 14241 |
| Rabbit-anti-TRAF6 | 1:1000 (WB) | Cell Signaling Technology | # 67591 |
| Mouse-anti-Myosin heavy chain | 1:500 (WB)/1:50 (IF) | DSHB | # MF20 |
| Mouse-anti-Myogenin | 1:500 (WB)/1:50 (IF) | Invitrogen | # MA5-11486 |
| Rabbit-anti-p-YAP1 | 1:1000 (WB) | Cell Signaling Technology | # 4911 |
| Rabbit-anti-YAP1 | 1:1000 (WB) | Cell Signaling Technology | # 14074 |
| Rabbit-anti-p-ERK1/2 | 1:1000 (WB) | Cell Signaling Technology | # 9101 |
| Rabbit-anti-ERK1/2 | 1:1000 (WB) | Cell Signaling Technology | # 9102 |
| Rabbit-anti-p-JNK1/2 | 1:1000 (WB) | Cell Signaling Technology | # 9251 |
| Rabbit-anti-JNK1/2 | 1:1000 (WB) | Cell Signaling Technology | # 9252 |
| Rabbit-anti-p-p38 | 1:1000 (WB) | Cell Signaling Technology | # 9211 |
| Rabbit-anti-p38 | 1:1000 (WB) | Cell Signaling Technology | # 9212 |
| Rabbit-anti-α-Tubulin | 1:1000 (WB) | Cell Signaling Technology | # 2144 |
| Rabbit-anti-GAPDH | 1:1000 (WB) | Cell Signaling Technology | # 2118 |
| Anti-Mouse IgG2b AF488 | 1:1000 (IF) | Invitrogen | # A21141 |
| Anti-Rabbit IgG | 1:2000 (WB) | Cell Signaling Technology | # 7074S |
| Anti-Mouse IgG | 1:2000 (WB) | Cell Signaling Technology | # 7076S |
